# Wild chimpanzee groups increase social connectivity prior to risky collective action

**DOI:** 10.64898/2026.06.15.732266

**Authors:** James Brooks, Roger Mundry, Catherine Crockford, Roman M. Wittig, Erin G. Wessling, Liran Samuni

## Abstract

Cooperation is foundational to complex sociality, yet presents profound evolutionary dilemmas – costs and benefits are rarely distributed evenly and the decision to collaborate or defect can involve a complex contextual calculus. These challenges are compounded when cooperation scales from pairs to groups. Group-level cooperation is fundamental to many species’ success, but how is it sustained and regulated in nature? One route to addressing this question is to examine how individuals reorganise affiliative interactions in anticipation of group-level cooperation. We examine such pre–cooperative reorganisation using long-term data (2013-2018) from three neighbouring groups of wild chimpanzees at Taï National Park, Côte d’Ivoire. We found that prior to risky proactive territorial defence, chimpanzees formed more broadly connected, yet more diffuse, affiliative networks. Specifically, chimpanzees groomed and played with more group members on days of proactive territorial defence, and this pattern was temporally-sensitive, with increased affiliation occurring before, not after, the cooperative act. Chimpanzees accessed a broader range of partners through increased interaction efficiency by switching between more partners with shorter interactions per partner. This pattern suggests a shared basis of dynamic social readjustment in advance of group-level social dilemmas in hominids, potentially providing foundations for formalized systems of affiliation in human societies.

## Introduction

From cells to societies, cooperation is a foundational evolutionary process, allowing collectives to overcome obstacles that individuals independently could not. As powerful as cooperation may be at promoting fitness to some, it is vulnerable to cheating and defection as it entails conferring fitness benefits to others, sometimes at a great personal cost [1]. In pairwise interactions, kinship, mutualism, and reciprocity can explain and predict many observed cooperative behaviours across taxa [2]. However, cooperation can also scale beyond dyads to include even entire groups, and with this increase in scale the potential opportunities for, costs, and benefits of both cooperation and defection can grow dramatically [3–5].

Human societies demonstrate a remarkable capacity to create and sustain group-level cooperative systems. From foraging parties to political parties, humans coalesce into overlapping and intersecting groups at all scales, enabling them to act together for collective benefits. The tendency to coalesce into groups is so strong and so fundamental to humans’ evolutionary success that “group-mindedness,” i.e., the cognitive process of representing oneself and others at the level of groups, is invoked as a central feature of human cognitive evolution [6]. The ability to represent and interact with groups underpins forms of cooperation that enable humans to thrive across diverse ecologies, from small-scale division of labour [7] to large-scale social institutions and cultural group norms [8]. Understanding the evolutionary roots of human-like group-mindedness and group-level cooperation are therefore fundamental to understanding human evolution itself.

Group-mindedness in humans and other great apes has been suggested to be, in part, an evolutionary adaptation to the cooperative context of intergroup conflict [6]. Because intergroup conflict is inherently higher-order (i.e., it cannot be reduced to a set of dyadic interactions [6]), strategies evolved in the context of purely dyadic cooperation in these conflicts are therefore unlikely to suffice to regulate and sustain cooperation. As such, the representation of oneself as part of a common group (i.e., group-mindedness), rather than as an independent entity, is suggested to promote the risky collective cooperation necessary to defend shared spaces under threat [6]. Collective cooperation in defending shared spaces is especially evolutionarily important as it allows groups to not only mitigate the direct costs of out-group threat, such as an individuals’ personal safety, but also to yield substantial fitness benefits by maintaining groups’ exclusive access to, and potentially expanding, these spaces [9,10]. In this context, parochial altruism theory suggests that human dispositions for ingroup collaboration and outgroup hostility are driven by a co-evolved mechanism [11]. But which proximate behaviors may have evolved to support the group-level cooperation problem that is maintaining these physical spaces? In other words, how do societies across species maintain individual participation in group-level cooperation?

One proposed mechanism of regulating participation in cooperation is affiliative contact between members of the group [12,13]. For example, African wild dogs (*Lycaon pictus*) show intense affiliative behaviour among groupmates, called “rallies”, prior to group hunting [14]. Similarly, affiliative contact such as rubbing and petting promotes longer cooperative displays in allied bottlenose dolphins (*Tursiops aduncus*; typically in the context of defending a female against other males [15]), while spotted hyenas’ (*Crocuta crocuta*) greeting behaviour is associated both with subsequent coalition participation [16] and predator mobbing [17]. Affiliation has also been associated with group-level cooperation when the stakes are high in the face of an out-group threat. For example, allopreening rate and duration in green woodhoopoes (*Phoeniculus purpureus*) increase in riskier parts of their territory [18], and in vervet monkeys (*Chlorocebus aethiops pygerythrus*) intermittent pauses during intergroup encounters are marked by a combination of aggression and grooming towards individuals who defected and cooperated, respectively, in the preceding bout [19]. However, we know much less about whether and how non-human animal groups may change affiliative behaviour in anticipation of risky, proactive group-level cooperation.

Wild chimpanzees (*Pan trogolodytes*) live in societies centered around shared spaces that they aggressively defend during border patrols and intergroup encounters, representing some of the most striking examples of proactive collective cooperation in a non-human species [20]. Here, collaboration underlies acts that have the ability to bring group-level benefits [21,22], but are simultaneously risky to undertake alone or when outnumbered in an “imbalance of power” [23,24]. The outcomes of this collective action problem, and whether these collaborations ultimately succeed or fail, depend on groupmates’ willingness to engage in these acts together [25,26]. Chimpanzee territorial defence presents a social dilemma whereby individuals face immediate and acute danger together with other members of their group, including those they only rarely interact or associate with, for the group to secure their space and reproductive output. Despite the evolutionary importance of maintaining their territories, limited work has investigated how chimpanzee groups scale their cooperation in response to this inherently higher-order top-down cooperation challenge.

We know that territoriality influences chimpanzees’ social behaviour in several ways. On days of intergroup competition and in their aftermath chimpanzee groups are more cohesive while exhibiting reduced within-group aggression [27]. Individual participation in territorial activity is more likely in the presence of kin and strong bond partners, as well as with increasing numbers of participants [25,26,28], and social play is associated with increased likelihood of play partners cooperating together [29]. Still, individual time budgets, and with them the depth and breadth of possible social investments [30], are limited. These limitations are especially pronounced when significant time and energy are dedicated to territorial defence [31]. Prior studies have focused primarily on how interactions and relationships between pairs of individuals influence rates of participation, with less attention to how the diversity of partners may be related to group-level cooperation. In the face of an out-group threat, how does chimpanzee group cooperation scale from close friendships into a united force?

Given the difficulties of regulating cooperation participation, affiliative interactions may offer support and provide mutual assurance before committing to risky cooperative acts. While affiliation supports cooperation in a range of species [14–19], including chimpanzees [29], it is relatively understudied how these affiliative acts scale and may be associated with cooperation at the group level. In the context of territorial defence, affiliating with a broader range of partners is likely to be particularly advantageous for broadcasting motivation and inducing the cooperation needed to achieve strength in numbers that can tip the balance between winning and losing.

In most species, affiliative behaviors like social grooming and play are typically limited to interactions of two individuals at a time. To expand the breadth of potential cooperation partners through dyadic interactions individuals therefore require more separate interactions, either by spending more total time socializing or by having shorter interactions. However, not all grooming and play are truly dyadic - higher-order forms of affiliation (i.e., affiliating with two or more individuals simultaneously) remain poorly understood (see [32–35]), but may provide an alternate means to reaching a greater number of potential partners when time is a finite resource [36]. In addition to simultaneous direct interactions, polyadic grooming also enables a unique mode of socialization otherwise not possible in dyadic settings - indirect partner access. Specifically, individuals may be part of the same grooming cluster (i.e., connected chains of direct grooming interactions) without directly grooming one another, but it is unclear whether this form of indirect interaction may serve distinct social functions. It remains speculative whether higher-order affiliation, through either direct or indirect interactions, may support the evolution and maintenance of higher-order cooperation. Girard-Buttoz et al. [37] reported frequent polyadic grooming in the chimpanzees of Taï National Park relative to the bonobos of Lui Kotale and suggested that this may be in part due to the greater out-group threat and group-level cooperation observed in chimpanzees, but it remains unexplored whether polyadic affiliation itself plays a role in supporting group-level cooperation through partner diversity access and efficiency on days of territorial defence against outgroups.

We therefore aim to elucidate the affiliative behavioural responses through which wild chimpanzees’ may scale and sustain group-level cooperation. Following the hypothesis that affiliative social interactions can promote cooperation and that the effectiveness of intergroup territorial defence increases with the number of participants, we predicted that (1) chimpanzees groom and play with a larger number of their group members during days of group defence, as well as access more partners *indirectly* through belonging to the same polyadic grooming cluster. Further, we (2) predicted that chimpanzees are more efficient in accessing partners during days of group defence, doing so by reaching a larger number of partners either faster, simultaneously (i.e., via polyadic interactions), or both. This expectation is grounded in the premise that activity budgets constrain the time available for social interactions, especially on days of intergroup conflict [31]. Increased partner counts on days of group defence itself alone, however, could be driven by either preparation for upcoming group conflict and/or a response to this stressful context upon returning to safety. Given the assumed importance of affiliation in supporting joint participation in risky cooperation, we therefore (3) predicted that the timing of increased affiliation partners occurs prior, but not subsequent, to the end of territorial activity.

These predictions hold true whether or not decisions are tailored to anticipated defensive activity or are consequences of some contextual factors that may independently increase social engagement. These predictions together would therefore suggest chimpanzee groups acutely adjust to contexts associated with territory defence by shifting their affiliation networks on these days to cover a wider set of potential affiliation partners, as opposed to selectively reinforcing few strong long-term bonds or simply responding to competition-induced stress. How affiliative patterns shift across these contexts in wild chimpanzees can help reveal the proximate behavioural mechanisms underlying their group-level cooperation, and as such, the evolutionary foundations that enable societies to coordinate and scale their collaboration in risky and uncertain collective action dilemmas.

## Results

### Unique Partners Model

We found that both male and female chimpanzees interacted affiliatively (i.e., groomed and played) with more unique partners on days when they participated in proactive group defence (females: odds ratio (OR): 1.83, 89% CI: [1.28, 2.78]; males: OR: 2.88, 89% CI: [1.69, 6.59]; Fig 1, Table 1) than control days, despite similar time spent in social investment (see supplementary materials). In comparison, hunting days were not associated with more unique partners in either sex relative to control days (all CIs cross 0; Table 1). These effects were estimated after accounting for seasonal fluctuations (sine and cosine terms of date), suggesting an effect of collective action context on social connectivity over and above ecological variation over the year. As expected from prior studies [38], we found that generally males interacted with more partners than females (at reference categories for variables interacting with sex [i.e. on control days] OR: 2.62, 89% CI: [2.19, 3.18]) and that both sexes interacted with more partners in the presence of maximally tumescent females (OR: 1.64, 89% CI: [1.44, 1.88]; Table 1).

**Fig 1:**
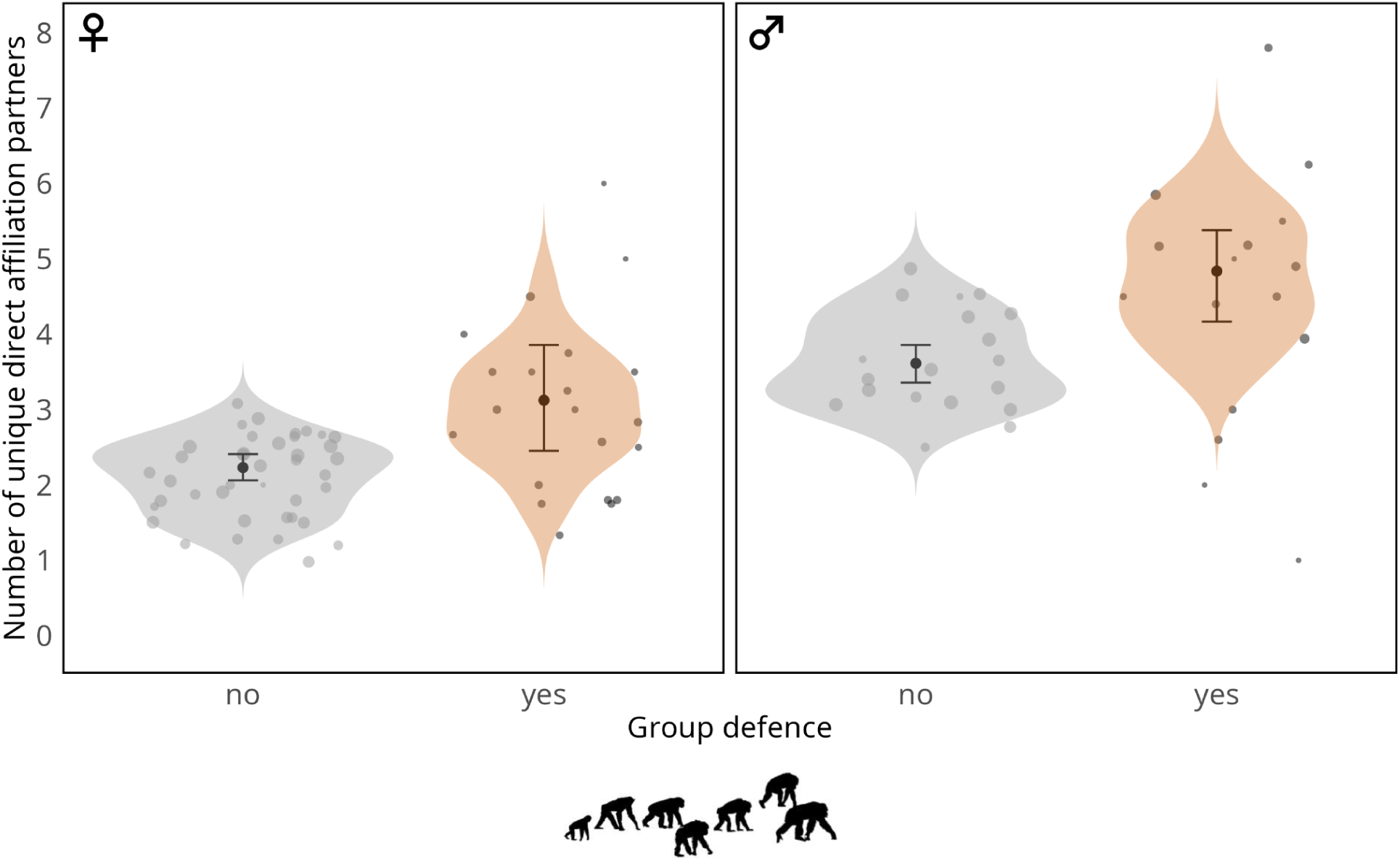
Increased partner connectivity in chimpanzees on days of group defence. Number of unique direct affiliation partners (grooming and play) by context (group defence vs no group defence) and sex (females on the left, males on the right). See supplementary material for modality-specific plots. Shown are the posterior means (black dots), 95% credible intervals (error bars), posterior distributions of the fitted (expected) probabilities from the model (violin), and individual play probabilities from the raw data (dots), with their area denoting the number of observations.

**Table 1:** Model estimates for the Unique Partners Model (direct grooming + play). Terms with an L indicate they make up the logistic component of the model.

| **parameter** | **Estimate** | **Error** | **95% CI** | **89% CI** |
| --- | --- | --- | --- | --- |
| Baseline (cLB) | -0.79 | 0.14 | -1.06, -0.52 | -1.01, -0.57 |
| Sex - Male (cLS) | 0.97 | 0.12 | 0.75, 1.21 | 0.78, 1.16 |
| Group defence (cLI) | 0.62 | 0.24 | 0.17, 1.14 | 0.25, 1.02 |
| Group defence × Sex (cLIS) | 0.50 | 0.46 | -0.32, 1.52 | -0.16, 1.27 |
| Hunt (cLH) | -0.05 | 0.15 | -0.34, 0.25 | -0.28, 0.19 |
| Hunt × Sex (cLHS) | -0.28 | 0.24 | -0.76, 0.20 | -0.67, 0.09 |
| Oestrus (cLO) | 0.50 | 0.08 | 0.34, 0.67 | 0.37, 0.63 |
| Date sine (cLSD) | 0.02 | 0.03 | -0.05, 0.09 | -0.03, 0.08 |
| Date cosine (cLCD) | 0.15 | 0.04 | 0.07, 0.24 | 0.09, 0.22 |
| Group - East (cLGE) | -0.36 | 0.12 | -0.61, -0.13 | -0.55, -0.17 |
| Group - North (cLGN) | 0.53 | 0.15 | 0.23, 0.83 | 0.29, 0.78 |
| Rate (cR) | -2.3 | 0.15 | -2.61, -2.01 | -2.55, -2.06 |
| Group size effect (cGS) | 3.47 | 0.16 | 3.20, 3.83 | 3.24, 3.75 |

These patterns (i.e., both sexes interacted with more partners on days of intergroup defence (but not hunts), males interacted with more unique partners than females overall, and individuals had more unique partners in the presence of maximally tumescent females) were found across both modalities of grooming and play (Fig S1, Tables S4, S5). Further, examining the impact of intergroup defence on the number of *indirect grooming* partners, that is individuals co-present in the same grooming cluster while not directly grooming one another, showed similar patterns (OR: 1.66, 89% CI: [1.03, 2.89]; Table S6). Specifically, individuals tended to be in grooming clusters with more unique partners on days of intergroup territorial activity across their interactions, although cluster sizes by bout were not on average larger than on control days (see Table S3).

### Efficiency Model

When modelling interaction efficiency by estimating the rate of partner accumulation over interaction time (i.e., how quickly chimpanzees reached new partners with increased total time spent affiliating), we found that polyadic grooming was more efficient than dyadic grooming (OR: 1.67, 89% CI: [1.54, 1.82]), and grooming was generally more efficient on days of group defence (OR: 1.49, 89% CI: [1.13, 2.04]; Fig 2, Table S11). However, there was no interaction between these effects (CI crosses 0; Table S11), meaning that both dyadic and polyadic grooming were more efficient on group defence days, with neither being disproportionately more efficient than the other relative to the control context. Males and females groomed with similar efficiency (CI crosses 0; Table S11), and there were no clear interactions between sex and grooming type nor collective action context (CIs cross 0; Table S11). This indicates that both sexes groomed with similar efficiency overall, and both increased their efficiency similarly via polyadic grooming and on days of intergroup defence (over and above seasonal variation, which captures contextual fluctuations known to affect interaction tendencies). As expected, the presence of an oestrous female in the group increased grooming access efficiency (OR: 1.21, 89% CI: [1.13, 1.30]; Table S11). We also found that indirect partner access was more efficient in males (OR: 1.41, 89% CI: [1.25, 1.60]) but this pattern was not affected by collective action context (CI crosses 0; Table S13). Similarly, polyadic play was more efficient than dyadic play but there was no effect of collective action context on play efficiency (CI crosses 0; Table S12), though this model had considerable uncertainty. In other words, while polyadic play was associated with more partners accessed for the same time investment compared to dyadic play, we did not find strong evidence that chimpanzees accumulated their partners faster on days of intergroup defence through either type of play.

**Fig 2:**
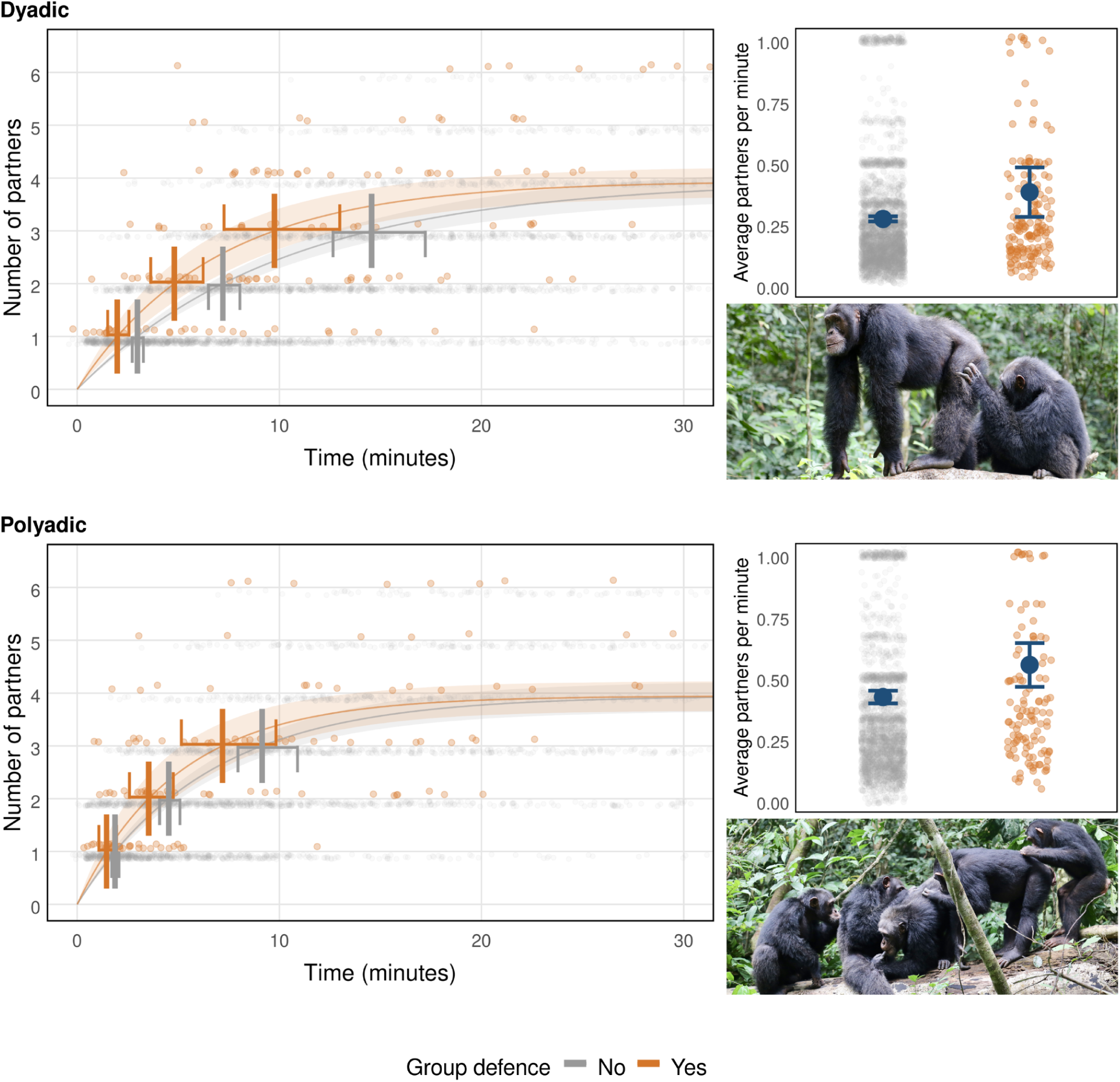
More efficient partner access via polyadic grooming and on days of group defence. Top: dyadic grooming partner accumulation over the first 30 minutes of grooming (left) and average rate in partners per minute (right). Bottom: polyadic grooming partner accumulation over the first 30 minutes of grooming (left) and average rate in partners per minute (right). All panels show results by context (group defence vs no group defence) Estimates and 95% credible intervals are model estimates over the dataset’s distribution of total interaction times. Photo credit: Liran Samuni, Taï Chimpanzee Project.

Timing of partner access relative to group defence activity:

We found that chimpanzees interacted with more unique partners before, but less-so after, group defence activity (Figs 3, Table S17; average time of group defence: M = 12:59, SD = 2.5h). We found this pattern to hold across all affiliation modalities investigated (all direct affiliation, grooming, play, and indirect grooming). More specifically, chimpanzees tended to interact affiliatively with an average of 1.5 partners both before and after control day pseudotimes and after group defence activity, while they interacted with more than 2.5 partners on average before group defence. Most of these affiliation partners were grooming partners. For play, chimpanzees on average interacted with around 0.1 partners before and after pseudotimes on control days and after actual group defence, but up to more than 0.5 partners on average before group defence. Finally, for indirect grooming, chimpanzees averaged around 0.5 partners for before/after pseudotimes on control days and following group defence, but up to 0.9 partners before actual group defence activity.

**Figure 3:**
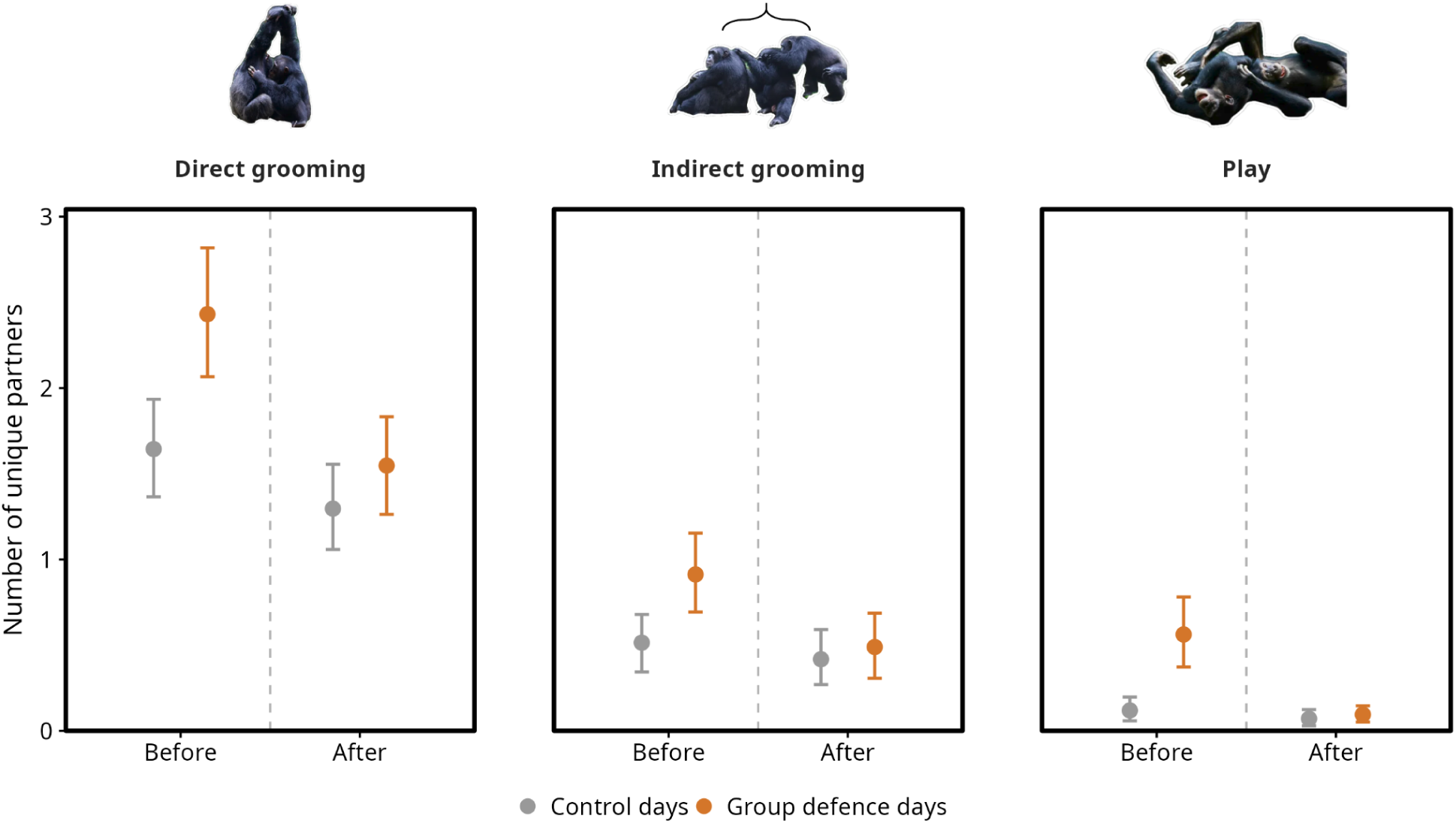
Increased partner access precedes group defence. Number of unique adult affiliation partners before and after times of group defence separated by interaction modality (left to right: direct grooming, indirect grooming [belonging to the same polyadic grooming cluster without direct contact], play). Points show the mean number of unique partners per observation across group defence days and 1000 resamples of an equal number of matched control days. Error bars are 95% intervals of the estimated mean: for group defence days, non-parametric bootstrap resamples (with replacement) of observations; for control days, across all 1000 iterations.

## Discussion

Chimpanzee groups dynamically adjusted the distribution of their social affiliation contacts in association with the acute top-down group cooperation challenge of maintaining their physical space. We found that on days of proactive territorial defence, both male and female chimpanzees interacted affiliatively with more unique members of their group before the territorial defense occurred. Chimpanzees diversified the breadth of their affiliative contacts, as opposed to only reinforcing support from their closest allies, despite limited time investment and tighter activity budgets on days of territorial defence (see supplementary materials). Increased partner diversity was found for both grooming and play, supporting the robustness of the effect in altering interaction patterns across distinct affiliative modalities. In the case of grooming, increases in partner diversity were achieved through increasing interaction efficiency, with individuals covering a wider proportion of available partners at the cost of a decreased investment per partner. Finally, timing sensitivity suggests that this increased partner diversity preceded but did not follow risky and stressful (see [39,40]) territorial defence. Regardless of whether these are intentional decisions made by individual chimpanzees, increased social connectivity may promote cooperation (and potentially buffer against defection) in this risky group-level social dilemma.

Our findings are consistent with prior reports in social species, from pack-hunting canids to cooperatively-breeding birds and alliance-forming marine mammals [15–19], which link affiliation to cooperation, but crucially emphasize affiliative diversity as opposed to affiliation rates themselves in potentially supporting risky group-level cooperation in chimpanzees. The present findings extend the scope of affiliative contact beyond deepening strong long-term ties or mere physiological regulation, to dynamically broadening short-term ties within communities in association with group-level cooperation. In chimpanzees, affiliative contact in play and grooming have been considered as joint actions in themselves [41], and affiliative contact via play was found to promote later co-participation in group-level collective action in the same dyads [29]. Joint actions may therefore foster interdependent chains of commitment dynamics and support joint participation in group-level cooperation [42,43]. Social connectivity does not arise passively but requires individuals to actively choose to engage with new partners. In other words, affiliative actions may help generate commitment, and affiliative diversity within groups may therefore help scale cooperation beyond dyads to interconnected groups.

Analysing temporal fluctuations in affiliative partner access, we showed that increased partner diversity occurred prior to the proactive intergroup activity, implying a preemptive response [44,45]. While other work has evaluated the consequences of intergroup encounters in the ‘common enemy effect’ [46–49], behavioural changes that precede between-group conflict are less studied (but see [50]). Although behavioural shifts in advance of intergroup territorial defence raise questions about the degree of planning by individuals, we cannot dissociate affiliative diversity before intergroup defence as an intended consequence, from affiliative partner diversity and subsequent intergroup defence both being driven by a common, non-cognitive, process. In any case, this leads to an interaction structure marked by wider affiliative diversity. Given theoretical work on affiliation’s role in commitment [51], the importance of affiliative ties in supporting human cooperation [52,53], and recent findings on affiliation’s possible role in joint commitment in great apes [29,54,55], these proximate structural shifts may functionally support short-term participation in group-level cooperation. This functional role is not dependent on, but certainly does not preclude, individual planning or intentionality. In this direction, recent theoretical work highlights that while across many species affiliative interactions may help support shared goals, and that this may scale to group-level cooperative contexts, humans may be unique in how they leverage these shared core mechanisms in ritualized and institutionalized systems [53,56].

Our findings cannot establish the direction of causality, but mainly reveal an association between affiliative diversity and consequent territorial defence. It is possible that individuals anticipate future collective action and adjust their behaviour, that engagement in territorial defence is facilitated in part by motivational states shaped by affiliative diversity (which may be driven by other, external factors), or, more likely, that these influences are mutually reinforcing rather than forming a simple unidirectional causal chain. Given the proactive nature of chimpanzee group defense [25,26,57], the rise in oxytocin in anticipation of proactive defence (see below; [58]), and evidence that affiliation promotes cooperation in this context [29] in ways that are vital for fitness [21,22], we consider it unlikely that the observed effects are merely byproducts of unaccounted for external factors. To differentiate mechanistic pathways to the emergence of group cooperation, experimental work may target the cognitive and motivational processes underlying decisions to both interact affiliatively with more partners (e.g., whether, relative to a control context, anticipated cooperation tasks induce increased affiliative diversity) and to cooperate in collective action (e.g., whether cooperation is more likely to succeed immediately following affiliative interactions). In any case, we observe affiliative diversity preceding upcoming group-level cooperation. Behavioural shifts in advance of territorial defence thus may provide a shared basis for the formalization of affiliative contact observed in humans prior to collective action.

Our findings also highlight polyadic affiliation as an efficient means by which to accumulate new partners, but do not uniquely emphasize the importance of polyadic interactions in intergroup defence contexts (either positively or negatively). Instead, our findings are most suggestive of a general increase in rates of partner switching, and scaling-up (partner wise) of all affiliative interaction forms, both dyadic and polyadic. Chimpanzees use more switching, as opposed to more total interaction time or more simultaneous partners, as the primary means by which they increase novel partner access on days of territorial defence. Girard-Buttoz et al. [37] did not find polyadic grooming to be more efficient than dyadic grooming in chimpanzees when comparing whether focal days with a greater proportion of polyadic grooming (out of all grooming that day) were associated with a greater number of cumulative partners that day. On the other hand, our results show that polyadic grooming does facilitate faster accumulation of direct partners in absolute time, compared to partner accumulation rates during dyadic grooming, but that polyadic grooming was not selectively used on days of group defence. Together, these results reinforce the potential of polyadic grooming to efficiently reach a broad range of partners with limited interaction time, but that chimpanzees do not leverage this efficiency selectively in reaching new partners on days of intergroup territorial defence.

Further, the role of *indirect* interaction partners accessed through polyadic grooming (i.e., belonging to the same cluster but not directly grooming one another) remains largely unexplored in the literature. We found evidence that chimpanzees increased indirect partner access on days of intergroup defence (and that these partners were preemptively reached) but it remains unclear whether this has biological significance in influencing subsequent behaviour (e.g., by inducing a “sense of togetherness” in shared experience [59]) and what mediates the distribution of polyadic interactions at either proximate or ultimate levels [34]. Our results highlight the potential functional benefits of polyadic interactions for temporal efficiency, while the absence of selective usage of direct polyadic grooming on days of territorial defence, when efficiency is especially important, hints at possible constraints. To this end, because individuals have more choice in who they groom as opposed to who grooms them, and because individuals can only directly groom one individual at a time, strategies towards increasing affiliation partners are expected to be expressed as both direct dyadic grooming partners and indirect polyadic partners, but less so as direct polyadic connections (where simultaneously reaching multiple direct partners inherently depends on both mine and others’ decision as it requires at least one other individual to be grooming oneself).

More broadly, our findings are largely consistent with and support recent theoretical work on parochial cooperation [11,60] and the evolution of group-mindedness [6]. Notably, where parochial cooperation theory highlights the ultimate and proximate association between ingroup prosociality and outgroup hostility [11], our findings emphasize the key role of not just the rate but the distribution of ingroup affiliative relationships. In a context like group defence in which strength in numbers is a crucial determinant of collective outcomes [22], reaching a broad range of partners is consistent with predictions from theory that these contexts exert distinct social pressures compared to average days. We observed wider but more diffusely distributed (i.e., across more partners but less investment per partner) affiliative contact to ingroup members in such contexts. The top-down cooperative challenge of territorial defence has been suggested as a major selection pressure on the evolution of group-mindedness [6], which may be facilitated in the short-term by partner breadth. Our results highlight potentially adaptive pathways linking affiliation to group-level action by showing that chimpanzee groups modulate their interaction patterns in association with a proximate group-level cooperative context. In line with theoretical work on human cooperative evolution, our findings provide evidence of continuity with extant species. As such, this work supports the view that human evolution likely acted on shared behavioural foundations, shaped by divergent socioecological priorities, as opposed to entirely *de novo* behavioural adaptations emerging after the split from our most recent common ancestor.

Mechanistically, in line with the proposed shared bases for group cooperation in humans and other species, the hormone neuropeptide oxytocin has been highlighted in group-level evolutionary physiology [61,61] as well as group-level and affiliative behaviour in other species. For example, oxytocin rises in advance of intergroup defence in wild chimpanzees [39,58], is suggested to act in a biobehavioural feedback loop with affiliation across species [62,63], and modulates group association dynamics to be wider but shallower in open-field experiments [64].

The mechanistic pathways by which group-level cooperation is achieved are yet to be fully resolved, but varied lines of evidence converge in linking intergroup conflict, group cooperation, and oxytocin reactivity. Converging lines of evidence may suggest functional links between affiliative contact diversity and the hormonal and behavioural consequences of these contacts stimulating participation in collective action.

While our work adds to growing evidence of flexible affiliative group dynamics in advance of intergroup territorial defence, it is not comprehensive and several open questions remain for future studies. Most centrally, comparisons with other field sites and species will be important to validate the generalizability of these findings. Different communities of wild chimpanzees differ greatly in the extent and intensity of their intergroup relations as well as group size [65,66], and it will therefore be valuable to assess both whether groups that experience greater outgroup threat show broader affiliative connectivity and whether day-level *shifts* to social investment budgets, relative to their baseline, are moderated by the intensity of outgroup threat. Specifically, we predict that both baseline social connectivity and shifts to social connectivity on days of intergroup defence will be positively associated with the intensity of outgroup threat. Further, several new questions on polyadic affiliation and higher-order cooperation are opened by this investigation. For example, there has been almost no prior attention towards the potential importance of *indirect* partner access in such polyadic grooming. Indirect connections within polyadic grooming clusters are often removed from analysis, and it remains unclear whether and how these connections influence subsequent cooperation. Whether polyadic grooming is employed differently by groups of different sizes and facing different degrees of intergroup competition is a crucial empirical question for cross-population comparisons, where we predict that polyadic grooming will be most frequent in large, tolerant, and cohesive groups. Finally, comparison with other species to place the present findings in a phylogenetic comparative context will provide valuable data to reconstruct the evolutionary processes which shaped adaptive social responses in proactive group-level cooperation.

## Conclusion

Societies persist through cooperation. The group-level cooperative challenge of maintaining shared spaces is a fundamental evolutionary process at the centre of group living and sociality. Here, we showed that chimpanzees, whose societies are characterized by flexible social association dynamics and strong out-group threat, broaden their affiliative network within their group before engaging in risky and proactive territorial defence. In doing so, we highlight interaction breadth and efficiency as a key mechanism by which cooperation scales from dyadic to group level. Cooperation is central to how all societies survive and thrive, and this cooperation is supported, in part, by interconnected and interdependent affiliative contacts and commitment. In group-level cooperation, a wider diversity of interaction partners can support the scaling of this affiliative contact from pairwise bonds into large-scale coordinated collective action.

## Methods

### Ethics statement

All methods used in this study were non-invasive and were approved by the Ministries of Research and Environment of Côte d’Ivoire, and Office Ivoirien des Parcs et Reserves. All aspects of the study comply with the ethics policy of the American Society of Primatologists.

### Field site and data collection

We studied three neighboring groups (i.e., North, South, and East) of wild chimpanzees (*P. t. verus*) in Taï National Park, Côte d’Ivoire from 2013 to 2018. Systematic long-term data were collected using standardized protocols of full-day nest-to-nest focal animal follows [67]. As follows are sometimes interrupted, we included only all focal observation days with at least four hours of observation time (approximately 50% of a full-day’s follow [mean daily follow of 8.8 hours in this dataset]). Focal data included the timing and partners involved in all grooming and play interactions as well as all changes in activity (such as movement and feeding behaviour), in addition to participation in proactive intergroup defence (i.e., initiated intergroup encounters and border patrols; definition in supplementary materials). We also documented participation in monkey hunting, a cooperative group activity in which success depends more on the quality (rather than quantity) of partners [68] as a comparative point to intergroup defence. From these data, we extracted all unique grooming and play partners, the form (i.e., dyadic versus polyadic) and the duration of each interaction with these partners, as well as the cumulative time spent engaging in grooming, play, and feeding behaviour throughout the day. On days of group defence, we noted for each affiliative interaction whether it took place before or after the end of the group defence activity. Unless otherwise specified, our final dataset included 59 focal individuals over 4483 observation days, with a median of 66 days per individual (range: 1-175).

### Study design

Our main questions were how chimpanzees distributed their time and affiliative interaction decisions on days of proactive intergroup territorial defence, hunting, and control days that lacked these cooperative behaviours? In other words, is group-level cooperation associated with shifts in the timing and extent to which wild chimpanzees expand the number of partners with whom they affiliatively interact? We modeled the total unique number of partners directly accessed overall and by each form of affiliation (grooming and play) per day. In addition, we modeled the number of partners *indirectly* accessed via polyadic grooming (i.e., belonging to the same polyadic cluster but not directly grooming one another). These measures correspond to the social network metric of strength centrality in the focal individual’s daily unweighted egocentric affiliation network. Under our hypothesis, we predicted intergroup defence contexts (in which chimpanzees depend crucially on strength in numbers) but not hunts (in which partner quality is thought to be more important than quantity) to be associated with an increase in unique partners relative to control days when neither behavior occurred.

Next, we asked *how* chimpanzees could proximately increase their affiliation partners via exactly three potential, non-mutually-exclusive, means: a) more total time devoted to affiliation, b) more rapid switching between interactions, or c) more simultaneous unique partners (i.e., through polyadic social interactions). We, therefore, modeled partner access *efficiency* - the rate at which chimpanzees acquire new partners over time spent in affiliative interactions. We model efficiency by affiliation type (dyadic versus polyadic) and context (intergroup territorial defence versus control) as well as their interaction. We analyzed the efficiency of partner access separately for time spent in grooming and play interactions, as well as the efficiency of indirect grooming partner access (over time spent in polyadic grooming). According to our primary hypothesis, we predicted that both polyadic affiliation (as compared to dyadic) and affiliation on days of intergroup defence (compared to control days) are associated with a more rapid accumulation of unique partners over time. Alternatively, if affiliation prior to group defence is driven primarily by reinforcement of long-term dyadic bonds, we would instead predict that chimpanzees prepare for intergroup conflict by investing relatively more of their social investment time in their strongest existing bonds, and thereby *decrease* the efficiency of acquiring new partners.

Finally, to compare whether increasing partner counts represent preparation for or reinforcement of the group cooperative context, we assessed the relative affiliation partner access before and after intergroup defence. We compared both the partners accessed before intergroup defence and partners accessed after intergroup defence to the number of partners accessed during matched control periods on days without intergroup defence. Together, these analyses aimed to probe whether and how chimpanzee groups selectively and dynamically shift their patterns of interaction to denser and/or broader coverage of potential affiliation partners in preparation for intergroup competition.

See supplementary material (Table S1) for an overview of all analysis methods.

### Models

We fitted all models in a bayesian framework using R (version 4.3.3) [69], with the function *brm* of the R package “brms” (version 2.23.0) [70]. For all models, we used four chains with 4000 iterations (of which 2000 were warmup) and the default package control structure. We modeled the means through which chimpanzees accumulated social interaction partners according to context (i.e. presence of group cooperation on that day). Neither time nor the number of available partners is expected to affect partner accumulation linearly: as individuals interact, the pool of uncontacted partners shrinks and the marginal increase in new partners per unit time declines, violating linearity assumptions. Therefore, we fitted custom non-linear functions to address our objectives (see below for implementation).

### Unique Partners Model - Effects of context on daily unique interaction partners

In our model examining the effects of context on the number of daily interaction partners (hereafter, ‘Unique Partners Model’) we estimated how the proportion of group members with whom an individual interacted (i.e., the proportion of unique partners among all group members) was affected by various fixed effects predictors. The model consisted of three multiplicative components, one (the ‘logistic’ component) with our fixed effects and two other components (both exponential) to control for the non-linear effects of observation duration and group size on the number of partners accessed.

With the logistic component we modelled how the proportion of unique interaction partners was affected by the collective action context, namely, the presence of intergroup defence (no versus yes) and/or a hunt (no versus yes). Furthermore, we also controlled for the presence of oestrous females and seasonal variation (represented by the sine and cosine of Julian date expressed in radians; [71,72]), which are known to impact social interaction patterns [27], alongside group identity and the sex of the focal individual. Because the effects of the context may differ between the sexes, we also included the interaction between the sex of the focal and intergroup encounter and hunt and modelled all these as fixed effects. The logistic part of the model was implemented as follows:

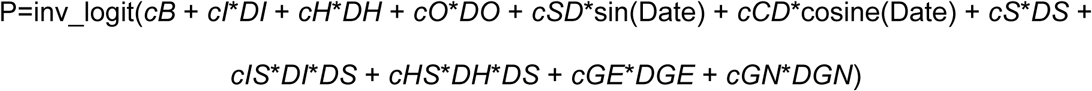

whereby *cB*, *cI*, etc. represent the estimated coefficients of the different terms (with *cB* as the intercept), *DI*, *DH*, etc. represent the dummy-coded predictors: *DI* = intergroup defence activity, *DH* = group hunting, *DO* = oestrous female, *DS* = focal’s sex (male), and *DGE* and *DGN* = group identity East and North, respectively, and *SD* and *CD* are the sine and cosine of date.

The inv_logit is a transformation with *x*’ 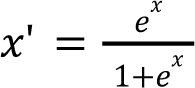 which ensures that fitted values are appropriately bound between zero and one, as required for proportion responses. In addition to the population-level model specified above, we also included varying effects (‘random intercepts’ and ‘random slopes’) for *cB, cI, cH, cO, cSD*, and *cCD* within the random effect of focal individuals. We *a priori* did not include parameters for the correlations among them to ease model fitting.

Naturally, the proportion of unique partners an individual interacted with increases over the cumulation of observation time in a day, for which we needed to control. Specifically, the proportion of unique partners can be expected to increase as an exponential function of observation time each day, which cannot be modelled as a standard linear term. Hence, we modelled this effect as follows:

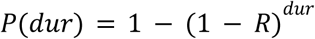

where *dur* is the observation duration, *R* is a parameter to be estimated, and P(*dur*) is the predicted proportion of interaction partners for this observation duration (measured in hours). To keep P(*dur*) bound between 0 and 1, we constrained *R* to the interval [0, 1] (see below for how we achieved parameter constraints). Note that P(*dur*) is zero when *dur* = 0, and approaches 1 as *dur* approaches infinity. This function therefore formed the second component of the model, which we thereafter refer to as the ‘exponential-of-duration’ component of the model.

Finally, because the proportion of unique interaction partners necessarily declines as group size increases, even if individual behaviour is otherwise identical, we controlled for group size. This term is the third component of the model (thereafter the ‘exponential-of-group-size’ term) implemented as:

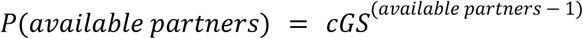

where *cGS* is a parameter to be estimated and was bound between 0 and 1. Note that P(available partners) = 1 when the group size is one (i.e., no available partners) and approaches 0 as group size approaches infinity.

We combined all three model components by multiplying them. Hence the fitted proportion was modelled as

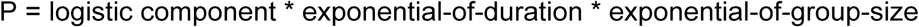

We fitted this model for four responses: a) the total unique direct affiliative interaction partners (i.e., both including play and direct grooming partners), b) unique direct grooming partners, c) unique play partners, and d) unique *indirect* grooming partners (i.e., individuals who at some point belonged to the same polyadic cluster while not directly grooming or being groomed by the focal individual).

### Efficiency Model - Effects of context on the efficiency of partner access

With a second group of models, we addressed how the efficiency of accessing partners was affected by intergroup defence context and interaction type (i.e., dyadic vs polyadic). Therefore, we modelled how unique partners accumulated over *interaction time*. As in the Unique Partners Model, the response in this model was effectively the proportion of group members an individual interacted with and the model again consisted of three parts - the logistic component, the exponential-of-duration component, and the exponential-of-group-size component – which were multiplied together. Here, however, the exponential-of-duration component estimated how the number of affiliative partners accumulated over increasing *time spent in affiliative interactions* as opposed to controlling for total observation time (as in the Unique Partners Model). We therefore modelled how the *rate* of partner accumulation was affected by group defence context (yes versus no) and interaction type (dyadic versus polyadic) and controlled for focal sex (and its interactions with defence context and interaction type) in the ‘exponential-of-duration’ component. As in the previous model, the ‘exponential-of-duration’ part was modelled as P(*dur*)=1-(1-*R*)^*dur*, but in this model, the term *R* was itself modelled as a linear combination of fixed effects, bound between zero and one, and *dur* represented *interaction* time (in minutes). Specifically, we modelled

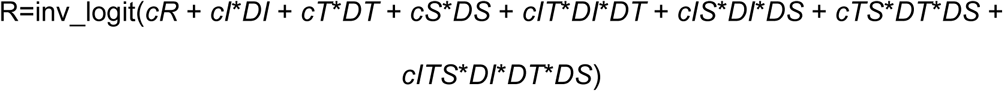

whereby *cI, cT*, etc. represent the estimated coefficients, *cR* is the intercept, and *DI, DT*, and *DS* are dummy variables for defence context, interaction type, and focal sex, respectively. Varying effects were included for *cI, cT*, and *cIT* within the random effect of focal individual.

We controlled for the presence of oestrous females, seasonal variation, and group identity in the ‘logistic component’ and the effect of group size in the ‘exponential of group size’ part as we did in the Unique Partners Model (see above). Specifically, for the logistic part we modelled

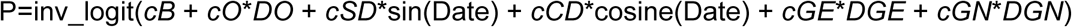

whereby *cB, cO*, etc. represent the estimated coefficients, *cB* is the intercept, and *DO*, *DGE*, and *DGN* represent the dummy predictors for presence of an oestrous female and group identities East and North, respectively (i.e., with South as the reference category). We again included varying effects here for *cB, cO, cSD*, and *cCD* by focal individual. Note that because a random intercept was included for *cB*, as in the first model, no random intercept was estimated in the exponential of duration component to avoid model redundancy.

Finally, the exponential-of-group-size component was identical to the previous model (i.e., implemented as *P*(*available partners*) = *cGS^(available partners − 1)^*). We again combined all three model components (i.e., exponential of duration part, logistic part, and exponential of group size part) by multiplying them.

With this model structure, we fitted separate models for grooming and play interactions, whereby *dur* represents the time observed engaged in the respective affiliation type only. For grooming, we additionally distinguished between direct grooming (i.e., individuals who groomed and/or received grooming from the focal) and indirect grooming (i.e., individuals who at some point belonged to the same polyadic cluster while not directly grooming or being groomed by the focal individual) in separate models. As indirect grooming only occurs as part of polyadic interactions, we did not include interaction type as a predictor in this model.

### Timing of partner access relative to group defence activity

We additionally analyzed how many partners were accessed before and after proactive intergroup defence, each compared to control days. To this end, we defined all partners accessed prior to the end of the group defence activity as “before” and all partners accessed following the end of the collective context as “after”. We aimed to compare these partner distributions on days of group defence to its expected distribution on control days. To achieve this, we ran 1000 iterations of randomly assigning real group defence times to control days and comparing the number of partners accessed before and after these times. More specifically, for each iteration, we first randomly sampled a set of control days equal in number to the number of intergroup defence days, additionally matched by group identity (see supplementary material for analysis further matching *focal* identity). Each of these control days was then assigned a “pseudotime” corresponding to the timing of group defence on its matched intergroup defence day. We ensured all group defence days in this analysis and all matched control days in all iterations had a minimum of one hour of observation both before and after the time of group defence. For each day, we then determined how many unique partners were accessed before and how many were accessed after this time or pseudotime. Once we had determined the number of partners relative to the time (or pseudotime) each day, we then established, for each matched pair of group defence and control days, the difference between the number of partners accessed (i.e., number of partners on group defence days minus number of partners on control days). A positive value represents, on average, more partners accessed on group defence days than control days, while a negative value represents more partners on control days. We calculated the difference in partners separately for the “before” and “after” periods for each matched pair, and averaged the differences across all pairs in this iteration. We repeated this process for 1000 iterations and then calculated the proportion of iterations whose average difference was smaller than or equal to zero as our test statistic. If this value is significant relative to chance (i.e. expecting roughly half of iterations to have more partners in group defence compared to control days), it implies the number of partners accessed was larger on days of group defence compared to control days. We also conducted the same analysis matching control days by both focal individual and group to verify the robustness of the result (see supplementary material).

All results in the main text show analyses focused only on individuals 11 years or older, but each of these analyses was additionally fitted for partners of all ages (see supplementary material).

We selected this age category for our main analyses because it is at this age that individuals are well-integrated into the dominance hierarchy, form strong and stable bonds (even reaching the alpha position for males), and regularly participate in intergroup territorial defence and hunting (see supplementary materials).

## Acknowledgements

We are grateful to the Ministère de l’Enseignement Supérieur et de la Recherche Scientifique (research permit: Wittig/006/MESRS/DGRI), the Ministère de Eaux et Forêts in Cote d’Ivoire, and the Office Ivoirien des Parcs et Reserves for allowing us to carry out this study. We thank the Centre Suisse de Recherches Scientifiques en Côte d’Ivoire for their logistical support, the Taï Chimpanzee Project research team for field assistance, and the Wild Chimpanzee Foundation, especially Christophe and Hedwige Boesch, for long-term commitment to the TCP.

