## supplementary materials for "Wild chimpanzee groups increase social connectivity prior to risky collective action"

### Supplementary Information

#### Supplementary methods:

Group defence definition: As group defense we only included instances of proactive territorial activity, that is border patrols and intergroup encounters that involved an approach behavior by the focal group. Border patrols are characterized by groups of chimpanzees quietly traveling towards or beyond their territorial borders, often involving periods of listening, smelling, and looking for signs of the neighbouring group [1,2]. Sometimes, but not always, these lead to an intergroup encounter, which can be vocal, visual, or physical interactions with individuals from another group. We included all border patrols in our definition of group defence activity, and all intergroup encounters during which the focal group actively moved towards the neighbouring group (i.e., excluding days on which the focal group was ambushed by another group). In the analyses, we noted the occurrence of group defence only when the focal individual participated in the territorial act (possible encounters and patrols that the focal individual did not join were not counted).

Hunting: We included all confirmed occurrences of monkey hunting (direct observation of hunting as well as meat consumption) within the focal party. In this population, the majority of hunting (86%) is done cooperatively with multiple individuals and individual hunt success rates are extremely low (16%) [3].

Age selection rationale: We considered all individuals aged 11+ to be “independent”. This cut-off was selected because it is at this age when chimpanzees in this population become fully integrated into the dominance hierarchy, participate regularly in intergroup defence, and make independent association and ranging decisions. Individuals as young as 12 have attained alpha status in this population, despite older competitors, and we therefore determined 11 as the threshold for being classified as “independent” in these analyses.

Model Implementation

We fitted all models with priors which we set as follows: for the non-linear effects of available partners (Unique Partner Model and Efficiency Model), observation duration (Unique Partners Model), and interaction duration (Efficiency Model), we first fitted a model considering only the nonlinear effects in a maximum likelihood framework. We then checked whether the fitted values seemed to be a reasonable fit for the observed response and then used the obtained parameters of the nonlinear function as the means of their respective priors. The standard deviations of these priors were set to 1. Hence these were moderately informative priors. For all other estimates, we set priors to a mean of 0 and a standard deviation of 2, and hence these were weakly informed priors. For the priors of the random effects (both intercept and slopes) we used the default distributions of brms. Starting values for each of the fixed effects parameters were always identical to the mean of the respective prior distribution. All bulk and tail ESSs across all models were over 1500 and Rhat values were 1.00 throughout.

We achieved parameter constraints not by bounding the space on which the function *brm* operated, but by internally transforming parameters from an unconstrained to an effective space. For instance, when an effective parameter needed to be bound between 0 and 1, we transformed the unconstrained parameter P with P'=exp(P) / (1+exp(P)), or when a parameter needed to be non-negative we transformed it as P'=exp(P). The response in all models was the number of unique interaction partners over all potentially available unique interaction partners (in *brms* syntax: n_unique_partner | trials(n_unique_available)). We fitted all models with binomial error structure and identity link function. We used the identity link function because the model construction ensured that fitted values were appropriately bound between 0 and 1. The final sample size in all models was 59 individuals over 4483 focal follow days.

Table S1: Overview of analyses. See full methods section for details of model components and implementation details.

| Analysis | Response | Test effect | Non-linear terms | Control terms | Responses |
|---|---|---|---|---|---|
| Unique Partners Model | Number of partners accessed during the day | Effect of presence of group defence and group hunting on expected partner access (including interactions with sex)<br><br>Terms are included in model's logistic component | -Observation time (exponential-of-duration component)<br><br>-Group size (exponential-of-group-size component) | Presence of oestrous females, sine/cosine of date (seasonality), group identity<br><br>The terms are included in the model's logistic component | All direct affiliation partners (groom + play)<br><br>Direct grooming partners<br><br>Play partners<br><br>Indirect grooming partners (belonging to same polyadic cluster but not directly grooming) |
| Efficiency Model | Number of partners accessed during the day | Effect of presence of group defence on *rate* of partner accumulation (including an interaction with sex)<br><br>Term is included in the model of exponential-of-duration component | -Interaction time (exponential-of-duration component)<br><br>-Group size (exponential-of-group-size component) | Presence of oestrous females, sine/cosine of date (seasonality), group identity<br><br>The terms are included in the model's logistic component | Direct grooming partners<br><br>Play partners<br><br>Indirect grooming partners |
| Timing sensitivity analysis | Number of partners accessed, separately for | Mean difference between partner counts on group | NA | Matched by focal group identity (main text) and by both focal | All direct affiliation partners (groom + play) |

| | | | | | |
|---|---|---|---|---|---|
| | before/after time of group defence | defence days and matched control days | | individual and group identity (supplementary material) | Direct grooming partners<br><br>Play partners<br><br>Indirect grooming partners |

**Supplementary results:**

Time budgets:

We found that on days of proactive intergroup territorial defence, chimpanzees spent an average of 182 minutes feeding and 24 minutes socializing with other independent individuals (> 11 years) through grooming (23 minutes) and play (1 minute), out of 33 minutes socializing with individuals across all ages through grooming (28 minutes) and play (5 minutes). In comparison, on control days chimpanzees spent an average of 243 minutes feeding and 23 minutes socializing with other independent individuals (grooming: 22 minutes, playing: 0.3 minutes) out of 33 minutes socializing with individuals across all ages (grooming: 31 minutes, play: 2 minutes). We did not compare feeding rates on days of group hunts due to hunting producing a slowly-processed food item itself and due to data collection procedures focusing on sharing events instead of feeding time following hunts.

Table S2: Polyadic grooming descriptives (of independent individuals). All numbers refer to interactions directly involving at least two independent individuals (i.e. the focal plus at least one independent-aged partner). Average cluster size was calculated by taking the weighted average of cluster sizes (both polyadic [>=3] and dyadic [=2]) by the time spent in interaction.  All probabilities and proportions reflect the average across one focal observation day.

| | Control days | Group defence days |
|---|---|---|
| Probability of any grooming | 0.83 | 0.92 |
| Probability of any play | 0.14 | 0.34 |
| Average time grooming | 22 minutes | 23 minutes |
| Average time playing | 0.3 minutes | 1 minute |

| | | |
|---|---|---|
| Probability of any polyadic grooming | 0.63 | 0.69 |
| Average polyadic grooming time | 8.02 minutes | 6.71 minutes |
| Proportion of grooming time that is polyadic | 0.33 | 0.27 |
| Average grooming cluster size (weighted) | 2.33 | 2.3 |
| Probability of any polyadic play | 0.0051 | 0.032 |
| Average polyadic playing time | 0.016 | 0.12 |
| Proportion of playing time that is polyadic | 0.022 | 0.052 |
| Average playing cluster size (weighted) | 2.05 | 2.12 |

Table S3: Polyadic grooming descriptives (all ages). Average cluster size was calculated by taking the weighted average of cluster sizes (both polyadic [>=3] and dyadic [=2]) by the time spent in interaction. All probabilities and proportions reflect the average across one focal observation day.

| | Control days | Group defence days |
|---|---|---|
| Probability of any grooming | 0.93 | 0.97 |
| Probability of any play | 0.41 | 0.65 |
| Average time grooming | 31 minutes | 28 minutes |
| Average time playing | 2 minutes | 5 minutes |
| Probability of any polyadic grooming | 0.70 | 0.66 |
| Average polyadic grooming time | 8 minutes | 7 minutes |
| Proportion of grooming time that is polyadic | 0.24 | 0.22 |
| Average grooming cluster size (weighted) | 2.4 | 2.3 |
| Probability of any polyadic play | 0.087 | 0.18 |
| Average polyadic playing time | 0.34 | 0.68 |
| Proportion of playing time that is polyadic | 0.094 | 0.090 |
| Average playing cluster size | 2.18 | 2.17 |

| |
|---|
| (weighted) |

Table S4: Model estimates for Unique Partners Model (independent individuals, direct grooming only). Estimates beginning with ‘cL’ denote terms present in the logistic model.

| parameter | Estimate | Error | 95% CI | 89% CI |
|---|---|---|---|---|
| Baseline (cLB) | -0.68 | 0.14 | -0.96, -0.41 | -0.91, -0.46 |
| Hunt (cLH) | -0.11 | 0.17 | -0.45, 0.23 | -0.38, 0.16 |
| Group defence (cLI) | 0.61 | 0.26 | 0.14, 1.18 | 0.22, 1.04 |
| Date sine (cLSD) | 0.01 | 0.04 | -0.07, 0.08 | -0.05, 0.06 |
| Date cosine (cLCD) | 0.11 | 0.04 | 0.03, 0.20 | 0.05, 0.19 |
| Oestrus (cLO) | 0.54 | 0.1 | 0.36, 0.75 | 0.39, 0.70 |
| Sex - Male (cLS) | 1.03 | 0.14 | 0.77, 1.31 | 0.82, 1.25 |
| Hunt × Sex (cLHS) | -0.27 | 0.28 | -0.82, 0.27 | -0.71, 0.17 |
| Group defence × Sex (cLIS) | 0.33 | 0.48 | -0.50, 1.34 | -0.34, 1.09 |
| Group - East (cLGE) | -0.42 | 0.13 | -0.68, -0.17 | -0.63, -0.22 |
| Group - North (cLGN) | 0.59 | 0.18 | 0.25, 0.95 | 0.31, 0.88 |
| Rate (cR) | -2.48 | 0.16 | -2.78, -2.16 | -2.73, -2.21 |
| Group size effect (cGS) | 3.48 | 0.17 | 3.20, 3.86 | 3.24, 3.78 |

Table S5: Model estimates for Unique Partners Model (independent individuals, play only), Estimates beginning with ‘cL’ denote terms present in the logistic model.

| parameter | Estimate | Error | 95% CI | 89% CI |
|---|---|---|---|---|
| Baseline (cLB) | -3.54 | 0.41 | -4.50, -2.87 | -4.24, -2.98 |
| Hunt (cLH) | 0.64 | 0.28 | 0.05, 1.17 | 0.17, 1.07 |
| Group defence (cLI) | 0.93 | 0.46 | -0.03, 1.79 | 0.18, 1.62 |
| Date sine (cLSD) | 0.14 | 0.08 | -0.02, 0.31 | 0.01, 0.28 |
| Date cosine (cLCD) | 0.96 | 0.11 | 0.75, 1.19 | 0.79, 1.14 |
| Oestrus (cLO) | 0.12 | 0.13 | -0.14, 0.37 | -0.09, 0.32 |
| Sex - Male (cLS) | 0.78 | 0.19 | 0.40, 1.17 | 0.47, 1.10 |
| Hunt × Sex (cLHS) | -0.48 | 0.36 | -1.17, 0.26 | -1.05, 0.12 |
| Group defence × Sex (cLIS) | 0.32 | 0.59 | -0.82, 1.57 | -0.58, 1.26 |
| Group - East (cLGE) | 0.44 | 0.22 | 0.01, 0.87 | 0.08, 0.79 |
| Group - North (cLGN) | 0.12 | 0.25 | -0.39, 0.62 | -0.29, 0.52 |
| Rate (cR) | -3.96 | 0.48 | -4.65, -2.75 | -4.56, -3.11 |
| Group size effect (cGS) | 4.41 | 0.68 | 3.21, 5.86 | 3.40, 5.58 |

Table S6: Model estimates for Unique Partners Model (independent individuals, indirect grooming). Estimates beginning with ‘cL’ denote terms present in the logistic model.

| parameter | Estimate | Error | 95% CI | 89% CI |
|---|---|---|---|---|
| Baseline (cLB) | -0.84 | 0.19 | -1.22, -0.46 | -1.14, -0.53 |
| Hunt (cLH) | -0.47 | 0.44 | -1.38, 0.38 | -1.19, 0.22 |
| Group defence (cLI) | 0.52 | 0.32 | -0.08, 1.19 | 0.03, 1.06 |
| Date sine (cLSD) | -0.26 | 0.07 | -0.40, -0.12 | -0.37, -0.14 |
| Date cosine (cLCD) | -0.01 | 0.09 | -0.19, 0.17 | -0.15, 0.14 |
| Oestrus (cLO) | 0.51 | 0.13 | 0.26, 0.77 | 0.30, 0.73 |
| Sex - Male (cLS) | 1.64 | 0.25 | 1.18, 2.18 | 1.25, 2.06 |
| Hunt × Sex (cLHS) | 0.02 | 0.71 | -1.35, 1.46 | -1.09, 1.15 |
| Group defence × Sex (cLIS) | 0.76 | 0.57 | -0.24, 1.98 | -0.07, 1.68 |
| Group - East (cLGE) | -1.45 | 0.25 | -1.97, -0.99 | -1.87, -1.07 |
| Group - North (cLGN) | 1.83 | 0.4 | 1.11, 2.68 | 1.23, 2.48 |
| Rate (cR) | -3.78 | 0.13 | -4.03, -3.52 | -3.98, -3.57 |
| Group size effect (cGS) | 3.59 | 0.28 | 3.15, 4.21 | 3.21, 4.07 |

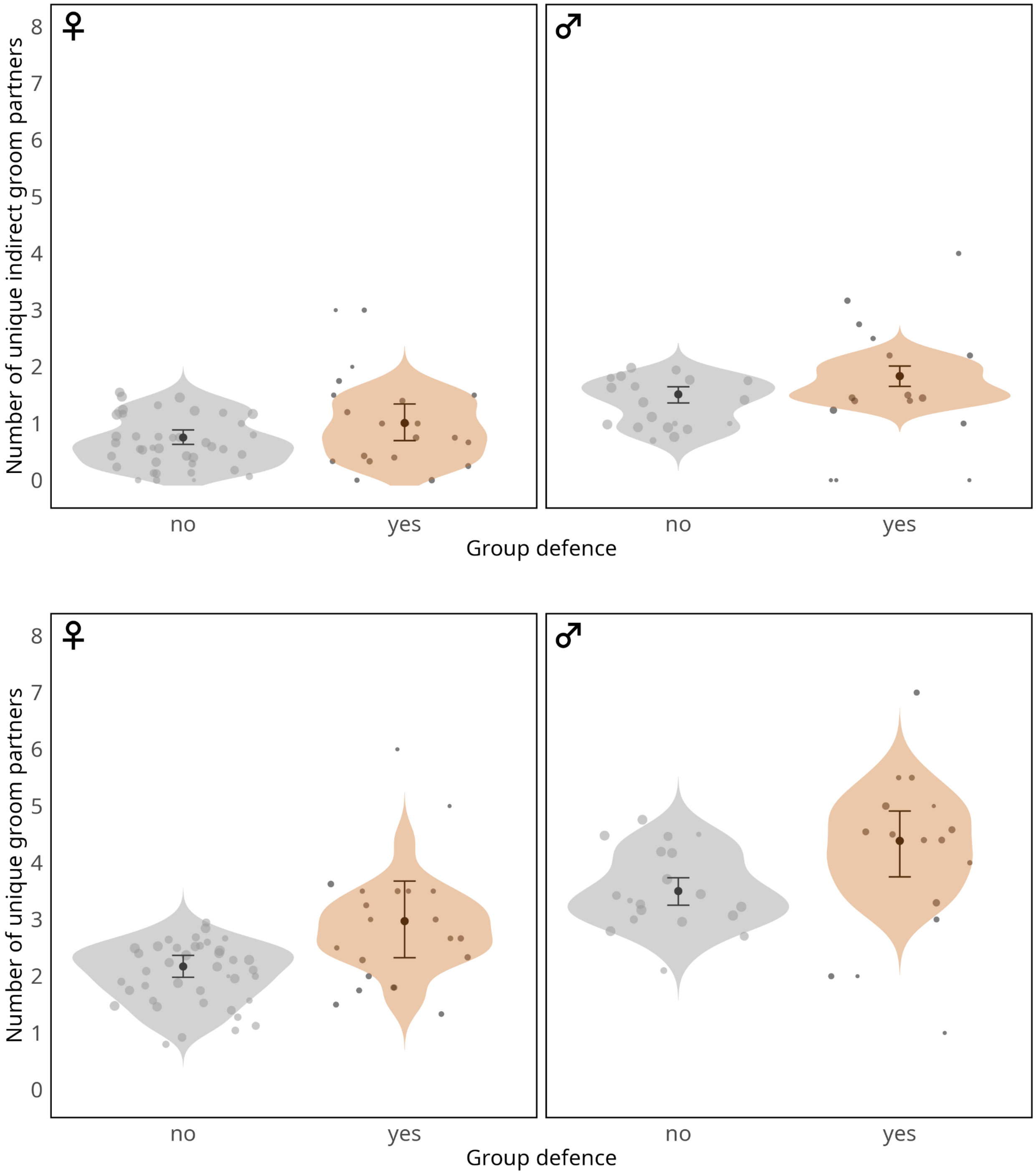

Number of unique indirect groom partners
Number of unique groom partners
0
1
2
3
4
5
6
7
8
♀
♂
no
yes
Group defence

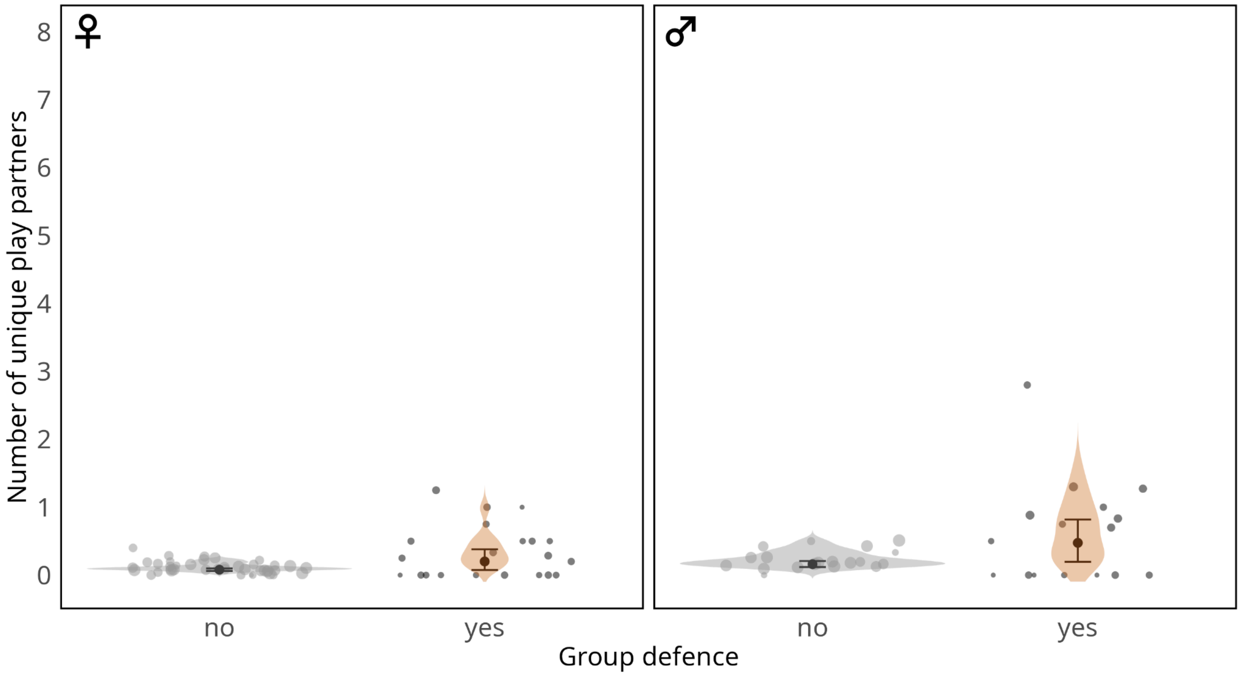

Figure S1: Number of unique affiliation partners (independent-age; top: indirect grooming, middle: direct grooming, bottom: play) by context (group defence vs no group defence) and sex (columns). Shown are the posterior means (black dots), 95% credible intervals (error bars), and posterior distributions of the fitted (expected) probabilities from the model (violin), and individual play probabilities from the raw data (dots), with the area denoting the number of observations.

Table S7: Model estimates for Unique Partners Model (all ages, all direct affiliation [grooming + play]). Estimates beginning with ‘cL’ denote terms present in the logistic model.

| parameter | Estimate | Error | 89% CI | 95% CI |
|---|---|---|---|---|
| Baseline (cLB) | -0.87 | 0.12 | -1.06, -0.68 | -1.10, -0.64 |
| Hunt (cLH) | -0.03 | 0.11 | -0.20, 0.15 | -0.25, 0.19 |
| Group defence (cLI) | 0.52 | 0.17 | 0.26, 0.80 | 0.21, 0.88 |
| Date sine (cLSD) | 0.04 | 0.02 | 0.00, 0.08 | -0.01, 0.09 |
| Date cosine (cLCD) | 0.16 | 0.03 | 0.11, 0.21 | 0.09, 0.22 |
| Oestrus (cLO) | 0.31 | 0.05 | 0.22, 0.39 | 0.21, 0.42 |
| Sex - Male (cLS) | 0.42 | 0.07 | 0.31, 0.53 | 0.28, 0.56 |
| Hunt × Sex (cLHS) | -0.30 | 0.17 | -0.59, -0.03 | -0.65, 0.03 |
| Group defence × Sex (cLIS) | 0.09 | 0.26 | -0.33, 0.51 | -0.43, 0.61 |
| Group - East (cLGE) | -0.22 | 0.08 | -0.35, -0.10 | -0.38, -0.07 |
| Group - North (cLGN) | 0.22 | 0.1 | 0.06, 0.37 | 0.03, 0.40 |
| Rate (cR) | -1.73 | 0.11 | -1.90, -1.56 | -1.94, -1.52 |
| Group size effect (cGS) | 3.66 | 0.08 | 3.53, 3.80 | 3.51, 3.84 |

Table S8: Model estimates for Unique Partners Model (all ages, direct grooming only). Estimates beginning with 'cL' denote terms present in the logistic model.

| parameter | Estimate | Error | 95% CI | 89% CI |
|---|---|---|---|---|
| Baseline (cLB) | -0.56 | 0.26 | -1.07, -0.07 | -0.99, -0.14 |
| Hunt (cLH) | -0.15 | 0.13 | -0.40, 0.10 | -0.35, 0.05 |
| Group defence (cLI) | 0.48 | 0.21 | 0.15, 0.95 | 0.20, 0.82 |
| Date sine (cLSD) | -0.01 | 0.03 | -0.07, 0.05 | -0.05, 0.04 |
| Date cosine (cLCD) | 0.05 | 0.03 | -0.02, 0.11 | 0.00, 0.10 |
| Oestrus (cLO) | 0.33 | 0.08 | 0.20, 0.50 | 0.22, 0.46 |
| Sex - Male (cLS) | 0.42 | 0.1 | 0.24, 0.65 | 0.27, 0.60 |
| Hunt × Sex (cLHS) | -0.22 | 0.2 | -0.63, 0.15 | -0.54, 0.08 |
| Group defence × Sex (cLIS) | 0.01 | 0.27 | -0.50, 0.59 | -0.40, 0.44 |
| Group - East (cLGE) | -0.37 | 0.11 | -0.60, -0.18 | -0.55, -0.21 |
| Group - North (cLGN) | 0.21 | 0.13 | -0.06, 0.47 | -0.01, 0.42 |
| Rate (cR) | -2.38 | 0.23 | -2.87, -1.99 | -2.77, -2.05 |
| Group size effect (cGS) | 3.73 | 0.14 | 3.49, 4.03 | 3.53, 3.97 |

Table S9: Model estimates for Unique Partners Model (all ages, play only). Estimates beginning with ‘cL’ denote terms present in the logistic model.

| parameter | Estimate | Error | 95% CI | 89% CI |
|---|---|---|---|---|
| Baseline (cLB) | -2.61 | 0.18 | -2.96, -2.27 | -2.89, -2.33 |
| Hunt (cLH) | 0.14 | 0.19 | -0.26, 0.49 | -0.17, 0.43 |
| Group defence (cLI) | 0.51 | 0.22 | 0.07, 0.95 | 0.15, 0.87 |
| Date sine (cLSD) | 0.15 | 0.04 | 0.09, 0.22 | 0.10, 0.21 |
| Date cosine (cLCD) | 0.62 | 0.07 | 0.47, 0.76 | 0.50, 0.74 |
| Oestrus (cLO) | 0.2 | 0.08 | 0.04, 0.35 | 0.07, 0.32 |
| Sex - Male (cLS) | 0 | 0.15 | -0.31, 0.31 | -0.25, 0.25 |
| Hunt × Sex (cLHS) | -0.23 | 0.27 | -0.76, 0.30 | -0.65, 0.20 |
| Group defence × Sex (cLIS) | 0.32 | 0.34 | -0.36, 0.96 | -0.23, 0.85 |
| Group - East (cLGE) | 0.27 | 0.17 | -0.07, 0.60 | 0.00, 0.53 |
| Group - North (cLGN) | 0.32 | 0.19 | -0.06, 0.70 | 0.01, 0.63 |
| Rate (cR) | -0.83 | 0.24 | -1.25, -0.29 | -1.19, -0.42 |
| Group size effect (cGS) | 3.04 | 0.08 | 2.91, 3.21 | 2.93, 3.17 |

Table S10: Model estimates for Unique Partners Model (all ages, indirect grooming). Estimates beginning with ‘cL’ denote terms present in the logistic model.

| parameter | Estimate | Error | 95% CI | 89% CI |
|---|---|---|---|---|
| Baseline (cLB) | -0.93 | 0.2 | -1.31, -0.52 | -1.24, -0.60 |
| Hunt (cLH) | -0.73 | 0.34 | -1.43, -0.10 | -1.27, -0.21 |
| Group defence (cLI) | 0.21 | 0.23 | -0.23, 0.66 | -0.16, 0.58 |
| Date sine (cLSD) | -0.25 | 0.06 | -0.37, -0.15 | -0.35, -0.17 |
| Date cosine (cLCD) | -0.07 | 0.08 | -0.23, 0.09 | -0.20, 0.06 |
| Oestrus (cLO) | 0.19 | 0.1 | 0.00, 0.38 | 0.04, 0.34 |
| Sex - Male (cLS) | 1.01 | 0.18 | 0.66, 1.39 | 0.73, 1.32 |
| Hunt × Sex (cLHS) | -0.13 | 0.5 | -1.12, 0.87 | -0.92, 0.65 |
| Group defence × Sex (cLIS) | 0.38 | 0.31 | -0.22, 0.99 | -0.10, 0.87 |
| Group - East (cLGE) | -1.32 | 0.2 | -1.74, -0.96 | -1.65, -1.02 |
| Group - North (cLGN) | 1.65 | 0.28 | 1.13, 2.23 | 1.22, 2.12 |
| Rate (cR) | -4.01 | 0.12 | -4.23, -3.76 | -4.19, -3.81 |
| Group size effect (cGS) | 4.74 | 0.46 | 4.02, 5.80 | 4.12, 5.56 |

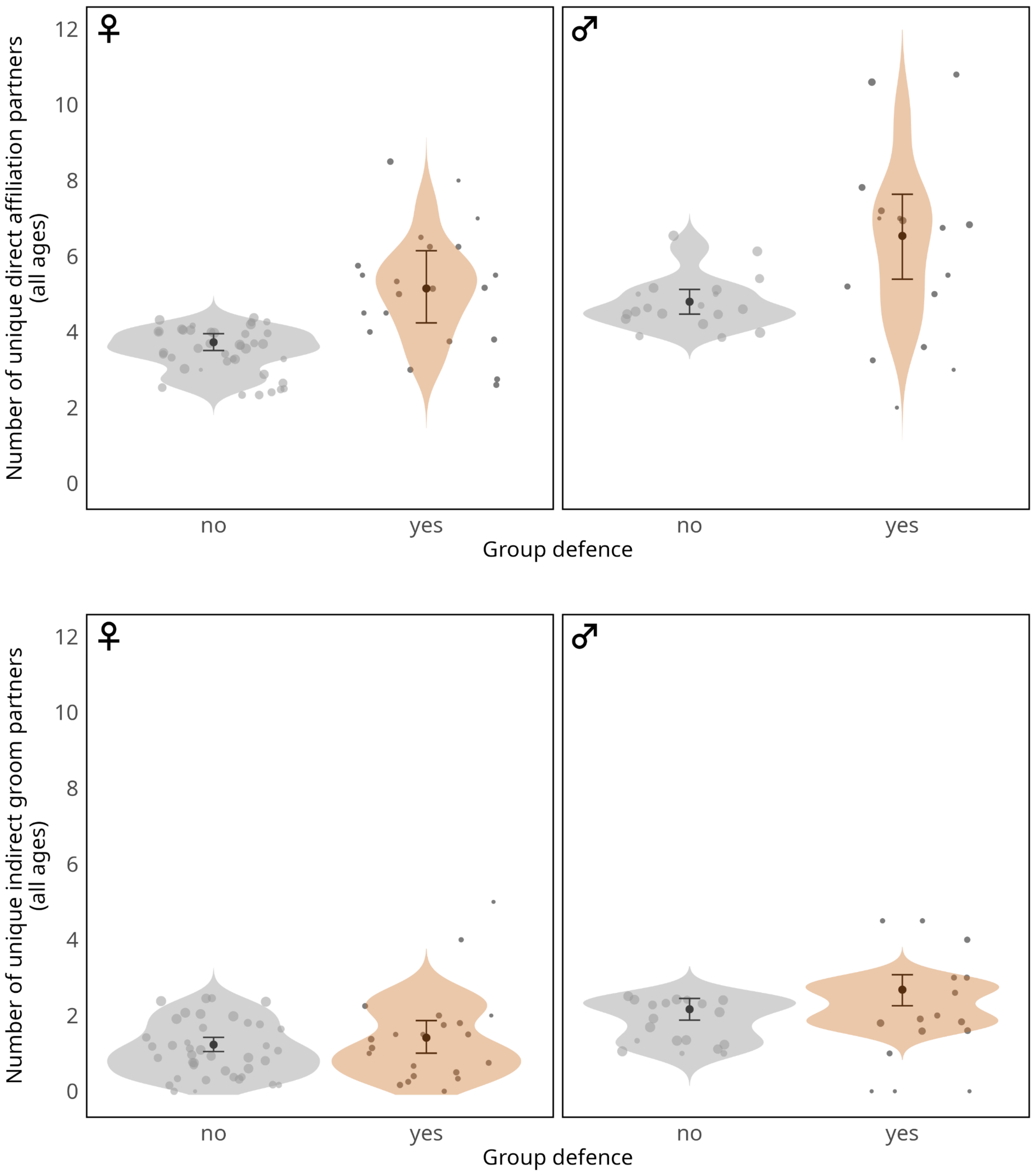
♀
♂
Number of unique direct affiliation partners
(all ages)
0
2
4
6
8
10
12
no
yes
no
yes
Group defence
♀
♂
Number of unique indirect groom partners
(all ages)
0
2
4
6
8
10
12
no
yes
no
yes
Group defence

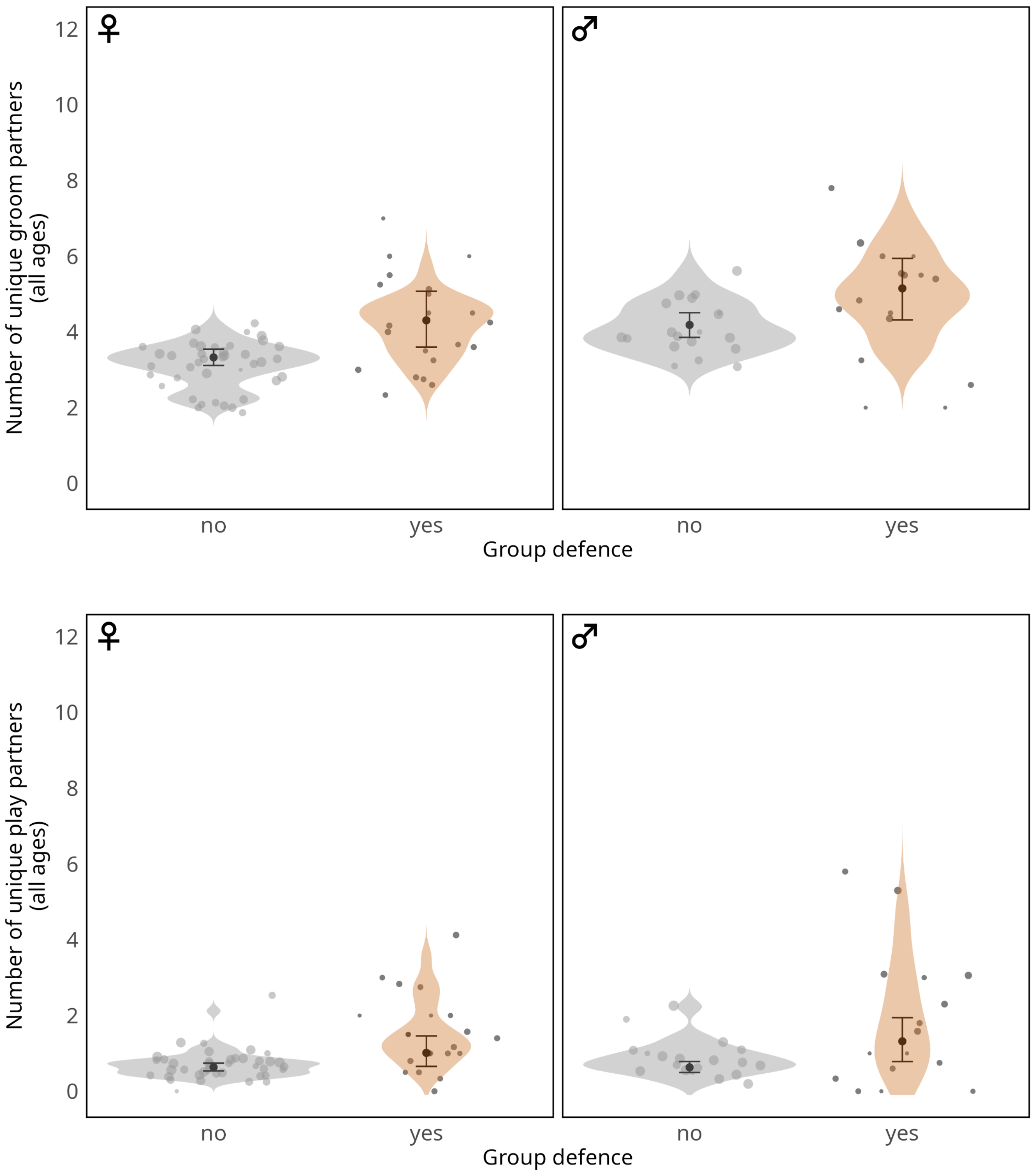

Figure S2: Number of unique affiliation partners (all ages; top to bottom: all direct affiliation, indirect grooming, direct grooming, play) by context (group defence vs no group defence) and sex. Shown are the posterior means (black dots), 95% credible intervals (error bars), and posterior distributions of the fitted (expected) probabilities from the model (violin), and individual play probabilities from the raw data (dots), with the area denoting the number of observations.

Table S11: Model estimates for direct grooming efficiency model (independent individuals). Estimates beginning with ‘cL’ denote terms present in the logistic model.

| parameter | Estimate | Error | 95% CI | 89% CI |
|---|---|---|---|---|
| Baseline (cLB) | -0.13 | 0.18 | -0.47, 0.24 | -0.41, 0.17 |
| Group defence effect on rate (cRI) | 0.41 | 0.19 | 0.06, 0.79 | 0.12, 0.71 |
| Affiliation type effect on rate (cRT) | 0.51 | 0.05 | 0.41, 0.62 | 0.43, 0.60 |
| Group defence × Affiliation type on rate (cRIT) | -0.24 | 0.28 | -0.78, 0.32 | -0.68, 0.22 |
| Date sine (cLSD) | 0.11 | 0.02 | 0.06, 0.16 | 0.07, 0.15 |
| Date cosine (cLCD) | 0.17 | 0.03 | 0.12, 0.22 | 0.13, 0.21 |
| Oestrus (cLO) | 0.19 | 0.04 | 0.11, 0.28 | 0.12, 0.26 |
| Base rate (cRB) | -2.26 | 0.05 | -2.37, -2.15 | -2.35, -2.17 |
| Sex effect on rate (cRS) | -0.03 | 0.07 | -0.17, 0.11 | -0.14, 0.08 |
| Group defence × Sex on rate (cRIS) | 0.06 | 0.26 | -0.44, 0.57 | -0.35, 0.48 |
| Sex × Affiliation type on rate (cRST) | -0.04 | 0.08 | -0.20, 0.11 | -0.17, 0.08 |
| Group defence × Sex × Affiliation type on rate (cRIST) | 0.2 | 0.38 | -0.56, 0.95 | -0.41, 0.80 |
| Group - East (cLGE) | 0.08 | 0.08 | -0.09, 0.24 | -0.05, 0.21 |
| Group - North (cLGN) | -0.01 | 0.11 | -0.24, 0.20 | -0.19, 0.17 |

| | | | | |
|---|---|---|---|---|
| Group size effect (cGS) | 3.08 | 0.11 | 2.89, 3.31 | 2.92, 3.26 |

Table S12: Model estimates for play efficiency model (independent individuals). Estimates beginning with 'cL' denote terms present in the logistic model.

| parameter | Estimate | Error | 95% CI | 89% CI |
|---|---|---|---|---|
| Baseline (cLB) | 0.75 | 1.16 | -1.61, 2.89 | -1.28, 2.53 |
| Group defence effect on rate (cRI) | 1.46 | 1.3 | -0.69, 4.38 | -0.35, 3.76 |
| Affiliation type effect on rate (cRT) | 2.69 | 1.5 | 0.02, 5.80 | 0.42, 5.23 |
| Group defence × Affiliation type on rate (cRIT) | 0.33 | 1.92 | -3.38, 4.11 | -2.66, 3.45 |
| Date sine (cLSD) | 0.08 | 0.14 | -0.21, 0.37 | -0.14, 0.30 |
| Date cosine (cLCD) | 0.29 | 0.15 | 0.00, 0.61 | 0.07, 0.54 |
| Oestrus (cLO) | 0.24 | 0.22 | -0.12, 0.74 | -0.06, 0.62 |
| Base rate (cRB) | 0.93 | 0.5 | 0.18, 2.12 | 0.28, 1.80 |
| Sex effect on rate (cRS) | 0.42 | 0.77 | -0.62, 2.55 | -0.44, 1.66 |
| Group defence × Sex on rate (cRIS) | 1.02 | 1.54 | -2.01, 4.16 | -1.37, 3.52 |
| Sex × Affiliation type on rate (cRST) | 1.77 | 1.68 | -1.47, 5.17 | -0.94, 4.49 |
| Group defence × Sex × Affiliation type on rate (cRIST) | 0.23 | 1.92 | -3.53, 4.06 | -2.82, 3.36 |
| Group - East (cLGE) | -0.59 | 0.56 | -1.91, 0.20 | -1.60, 0.12 |
| Group - North (cLGN) | -1.41 | 0.87 | -3.19, 0.10 | -2.86, -0.06 |

| | | | | |
|---|---|---|---|---|
| Group size effect (cGS) | 2.24 | 0.3 | 1.98, 3.14 | 2.00, 2.84 |

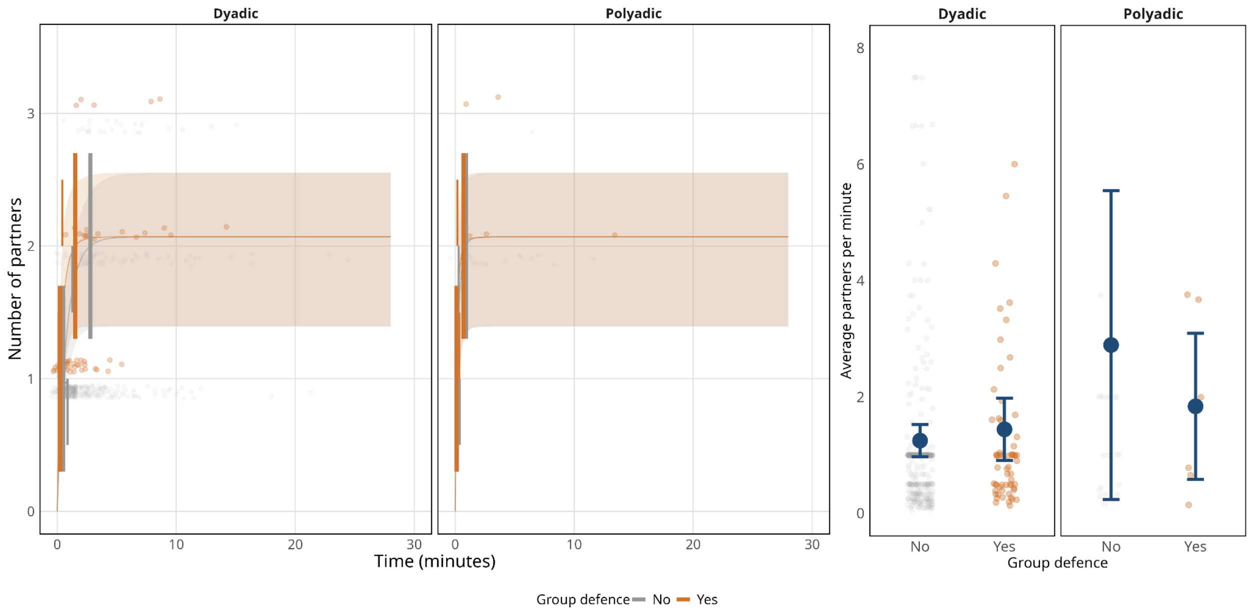

Figure S3: Dyadic (left) and polyadic (right) play partner (independent-aged) accumulation over the first 30 minutes of playing (left) and average rate in partners per minute (right). Estimates and 95% credible intervals are model estimates over the dataset's distribution of total interaction times.

Table S13: Model estimates for indirect grooming efficiency model (independent individuals). Estimates beginning with ‘cL’ denote terms present in the logistic model.

| parameter | Estimate | Error | 95% CI | 89% CI |
|---|---|---|---|---|
| Baseline (cLB) | 0.44 | 0.39 | -0.29, 1.23 | -0.16, 1.08 |
| Group defence effect on rate (cRI) | 0.23 | 0.19 | -0.13, 0.61 | -0.06, 0.54 |
| Date sine (cLSD) | 0.2 | 0.06 | 0.09, 0.34 | 0.11, 0.31 |
| Date cosine (cLCD) | 0.28 | 0.07 | 0.16, 0.43 | 0.18, 0.39 |
| Oestrus (cLO) | 0.21 | 0.11 | 0.04, 0.46 | 0.06, 0.40 |
| Base rate (cRB) | -2.93 | 0.08 | -3.08, -2.77 | -3.05, -2.80 |
| Sex effect on rate (cRS) | 0.34 | 0.08 | 0.19, 0.50 | 0.22, 0.47 |
| Group defence × Sex on rate (cRIS) | 0.28 | 0.25 | -0.21, 0.77 | -0.11, 0.68 |
| Group - East (cLGE) | -0.26 | 0.21 | -0.71, 0.09 | -0.62, 0.03 |
| Group - North (cLGN) | -0.16 | 0.23 | -0.66, 0.24 | -0.54, 0.18 |
| Group size effect (cGS) | 2.83 | 0.14 | 2.61, 3.15 | 2.64, 3.07 |

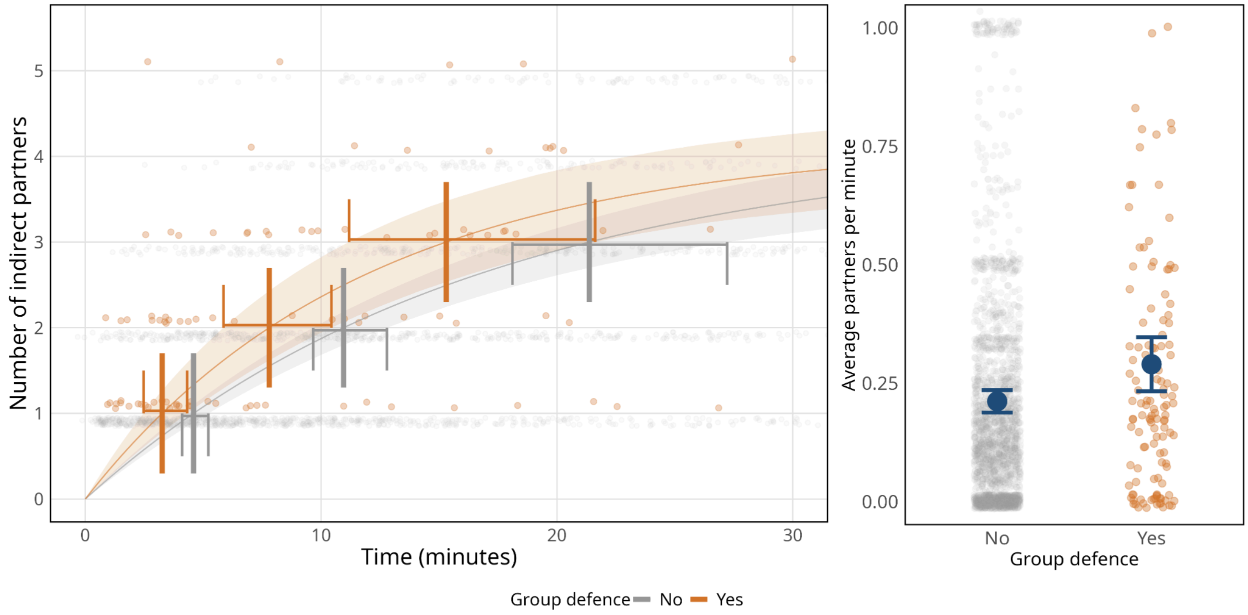

Figure S4: Indirect grooming partner (independent-aged) accumulation over the first 30 minutes of playing (left) and average rate in partners per minute (right). Estimates and 95% credible intervals are model estimates over the dataset's distribution of total interaction times.

Table S14: Model estimates for direct grooming efficiency model (all ages). Estimates beginning with ‘cL’ denote terms present in the logistic model.

| parameter | Estimate | Error | 95% CI | 89% CI |
|---|---|---|---|---|
| Baseline (cLB) | -0.55 | 0.17 | -0.87, -0.20 | -0.81, -0.28 |
| Group defence effect on rate (cRI) | 0.27 | 0.15 | -0.02, 0.58 | 0.03, 0.52 |
| Affiliation type effect on rate (cRT) | 1.15 | 0.05 | 1.05, 1.25 | 1.07, 1.23 |
| Group defence × Affiliation type on rate (cRIT) | -0.01 | 0.27 | -0.52, 0.56 | -0.43, 0.44 |
| Date sine (cLSD) | 0.06 | 0.02 | 0.02, 0.09 | 0.03, 0.08 |
| Date cosine (cLCD) | 0.12 | 0.02 | 0.08, 0.15 | 0.09, 0.15 |
| Oestrus (cLO) | 0.17 | 0.03 | 0.11, 0.24 | 0.12, 0.22 |
| Base rate (cRB) | -2.65 | 0.05 | -2.73, -2.56 | -2.72, -2.57 |
| Sex effect on rate (cRS) | 0.08 | 0.06 | -0.05, 0.20 | -0.03, 0.18 |
| Group defence × Sex on rate (cRIS) | 0.2 | 0.23 | -0.23, 0.65 | -0.15, 0.57 |
| Sex × Affiliation type on rate (cRST) | -0.51 | 0.07 | -0.65, -0.37 | -0.62, -0.39 |
| Group defence × Sex × Affiliation type on rate (cRIST) | -0.02 | 0.36 | -0.73, 0.66 | -0.60, 0.53 |
| Group - East (cLGE) | 0.05 | 0.07 | -0.09, 0.17 | -0.06, 0.15 |
| Group - North (cLGN) | -0.04 | 0.1 | -0.24, 0.14 | -0.21, 0.11 |

| | | | | |
|---|---|---|---|---|
| Group size effect (cGS) | 3.47 | 0.09 | 3.30, 3.67 | 3.33, 3.63 |

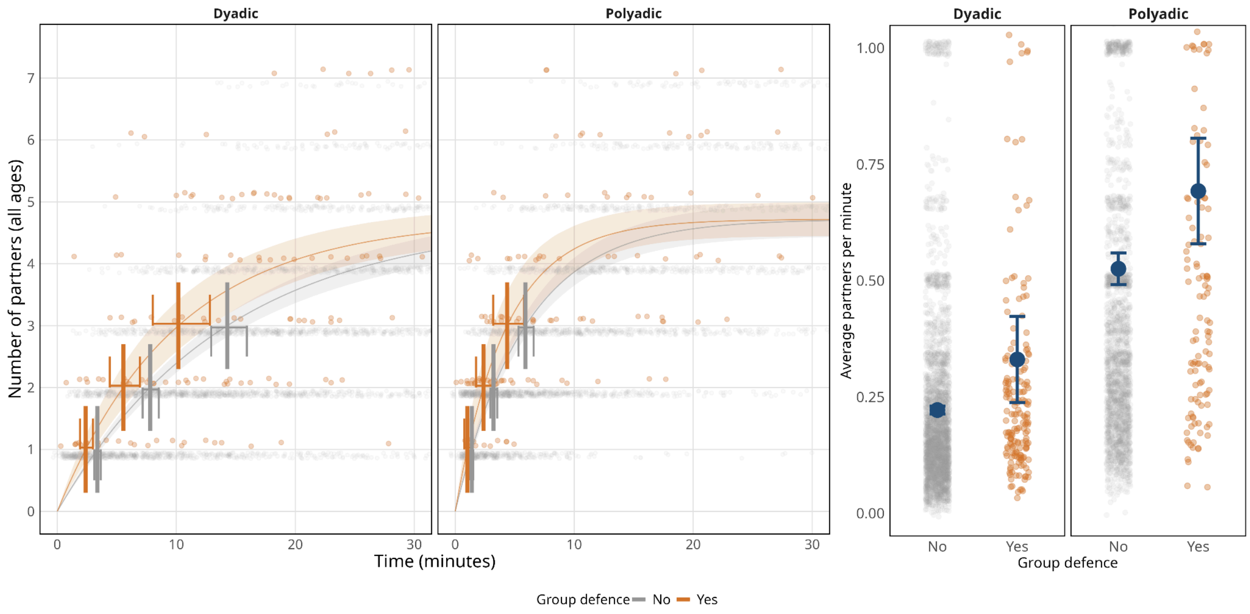

Figure S5: Dyadic (left) and polyadic (right) grooming partner (all ages) accumulation over the first 30 minutes of playing (left) and average rate in partners per minute (right). Estimates and 95% credible intervals are model estimates over the dataset's distribution of total interaction times.

Table S15: Model estimates for play efficiency model (all ages). Estimates beginning with ‘cL’ denote terms present in the logistic model.

| parameter | Estimate | Error | 95% CI | 89% CI |
|---|---|---|---|---|
| Baseline (cLB) | -1.08 | 0.32 | -1.69, -0.43 | -1.58, -0.55 |
| Group defence effect on rate (cRI) | 0.21 | 0.42 | -0.48, 1.17 | -0.38, 0.93 |
| Affiliation type effect on rate (cRT) | 5.02 | 0.97 | 3.38, 7.14 | 3.63, 6.71 |
| Group defence × Affiliation type on rate (cRIT) | 1.18 | 1.66 | -1.97, 4.58 | -1.42, 3.94 |
| Date sine (cLSD) | -0.01 | 0.03 | -0.08, 0.05 | -0.06, 0.04 |
| Date cosine (cLCD) | 0.17 | 0.05 | 0.08, 0.26 | 0.10, 0.25 |
| Oestrus (cLO) | 0.07 | 0.05 | -0.03, 0.17 | -0.01, 0.15 |
| Base rate (cRB) | -0.97 | 0.13 | -1.20, -0.71 | -1.16, -0.76 |
| Sex effect on rate (cRS) | 0.47 | 0.15 | 0.16, 0.76 | 0.22, 0.71 |
| Group defence × Sex on rate (cRIS) | 0.63 | 0.58 | -0.51, 1.83 | -0.29, 1.56 |
| Sex × Affiliation type on rate (cRST) | 0.98 | 1.53 | -1.76, 4.20 | -1.33, 3.53 |
| Group defence × Sex × Affiliation type on rate (cRIST) | 0.56 | 1.85 | -3.09, 4.32 | -2.37, 3.59 |
| Group - East (cLGE) | 0.12 | 0.08 | -0.03, 0.27 | 0.00, 0.24 |
| Group - North (cLGN) | -0.37 | 0.16 | -0.71, -0.07 | -0.64, -0.12 |

| | | | | |
|---|---|---|---|---|
| Group size effect (cGS) | 3.29 | 0.19 | 2.99, 3.71 | 3.03, 3.61 |

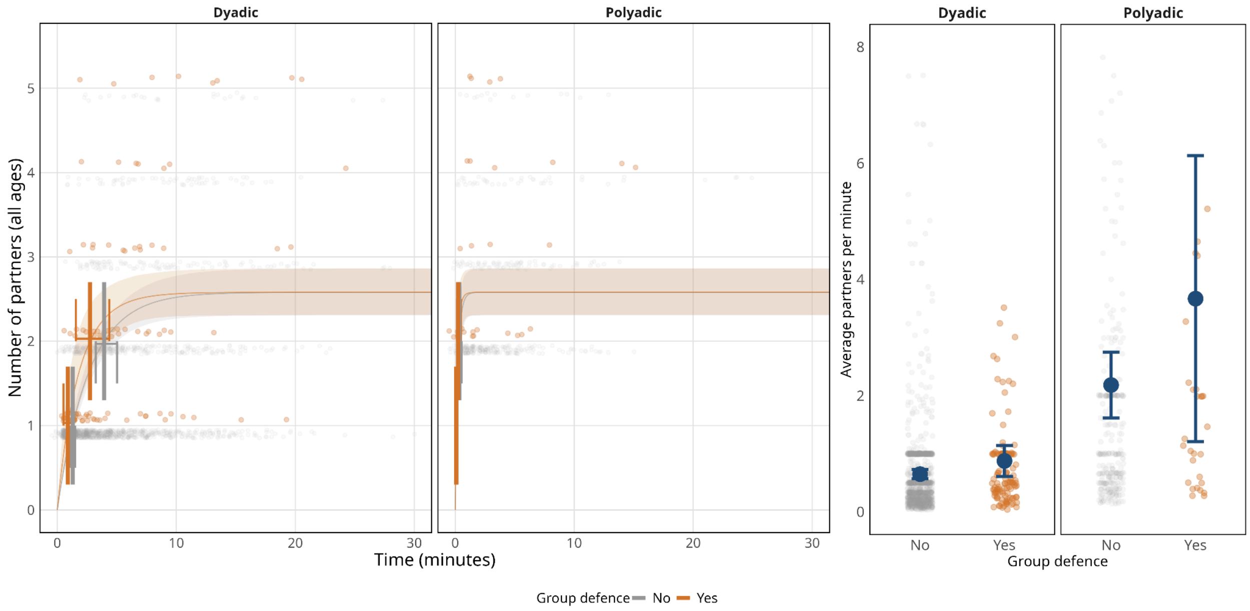

Figure S6: Figure X: Dyadic (left) and polyadic (right) play partner (all ages) accumulation over the first 30 minutes of playing (left) and average rate in partners per minute (right). Estimates and 95% credible intervals are model estimates over the dataset's distribution of total interaction times.

Table S16: Model estimates for indirect grooming efficiency model (all ages). Estimates beginning with ‘cL’ denote terms present in the logistic model.

| parameter | Estimate | Error | 95% CI | 89% CI |
|---|---|---|---|---|
| Baseline (cLB) | 0.18 | 0.4 | -0.54, 1.01 | -0.43, 0.83 |
| Group defence effect on rate (cRI) | 0.12 | 0.17 | -0.20, 0.46 | -0.15, 0.39 |
| Date sine (cLSD) | 0.09 | 0.04 | 0.01, 0.18 | 0.02, 0.16 |
| Date cosine (cLCD) | 0.18 | 0.06 | 0.08, 0.33 | 0.10, 0.29 |
| Oestrus (cLO) | 0.1 | 0.07 | -0.02, 0.24 | 0.00, 0.21 |
| Base rate (cRB) | -3.03 | 0.07 | -3.17, -2.89 | -3.14, -2.91 |
| Sex effect on rate (cRS) | 0.27 | 0.08 | 0.12, 0.42 | 0.15, 0.39 |
| Group defence × Sex on rate (cRIS) | 0.32 | 0.23 | -0.14, 0.79 | -0.04, 0.69 |
| Group - East (cLGE) | -0.29 | 0.16 | -0.65, -0.01 | -0.57, -0.06 |
| Group - North (cLGN) | -0.04 | 0.21 | -0.52, 0.32 | -0.41, 0.26 |
| Group size effect (cGS) | 3.4 | 0.15 | 3.16, 3.73 | 3.19, 3.66 |

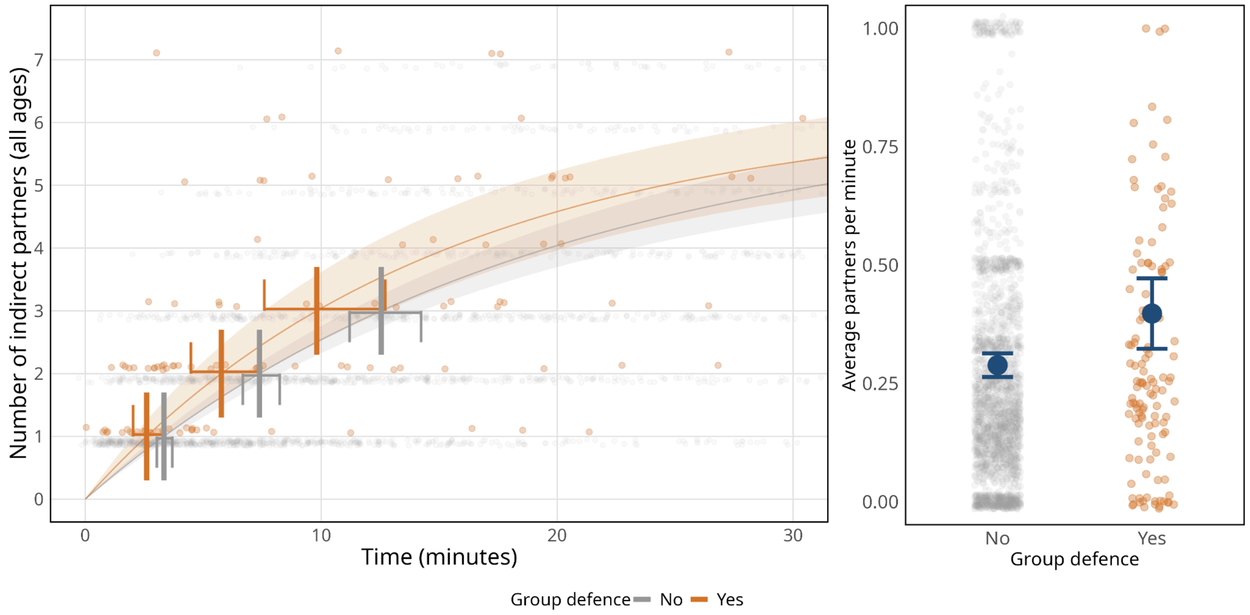

Figure S7: Indirect grooming partner (all ages) accumulation over the first 30 minutes of playing (left) and average rate in partners per minute (right). Estimates and 95% credible intervals are model estimates over the dataset's distribution of total interaction times.

**Timing of partner access relative to group defence activity (matched by group):**

**Independent individuals:**

Table S17: Estimated number of partners by affiliation modality for independent partners.

| Behaviour | Period | Days of group defence | Control days | Mean difference [95% quantiles] | p |
|---|---|---|---|---|---|
| Direct affiliation | Before | 2.78 | 1.72 | 1.06 [0.77, 1.34] | <0.001 |
| Direct affiliation | After | 1.64 | 1.34 | 0.29 [0.04, 0.53] | 0.034 |
| Direct grooming | Before | 2.43 | 1.64 | 0.79 [0.50, 1.07] | <0.001 |
| Direct grooming | After | 1.55 | 1.3 | 0.25 [-0.01, 0.49] | 0.064 |
| Play | Before | 0.56 | 0.12 | 0.44 [0.36, 0.50] | <0.001 |
| Play | After | 0.09 | 0.07 | 0.02 [-0.03, 0.07] | 0.434 |
| Indirect grooming | Before | 0.91 | 0.51 | 0.40 [0.23, 0.57] | <0.001 |
| Indirect grooming | After | 0.49 | 0.42 | 0.07 [-0.10, 0.22] | 0.376 |

**All ages:**

Table S18: Estimated number of partners by affiliation modality for all partners.

| Behaviour | Period | Days of group defence | Control days | Mean difference [95% quantiles] | p |
|---|---|---|---|---|---|
| Direct affiliation | Before | 4.5 | 2.67 | 1.82 [1.42, 2.18] | <0.001 |
| Direct affiliation | After | 2.37 | 2.06 | 0.31 [-0.03, 0.61] | 0.082 |
| Direct grooming | Before | 3.17 | 2.3 | 0.87 [0.52, 1.18] | <0.001 |
| Direct grooming | After | 2.04 | 1.8 | 0.24 [-0.07, 0.54] | 0.136 |
| Play | Before | 1.69 | 0.54 | 1.15 [0.95, 1.32] | <0.001 |
| Play | After | 0.45 | 0.37 | 0.08 [-0.07, 0.21] | 0.296 |
| Indirect grooming | Before | 1.17 | 0.76 | 0.41 [0.18, 0.63] | 0.002 |
| Indirect grooming | After | 0.63 | 0.62 | 0.01 [-0.22, 0.21] | 0.986 |

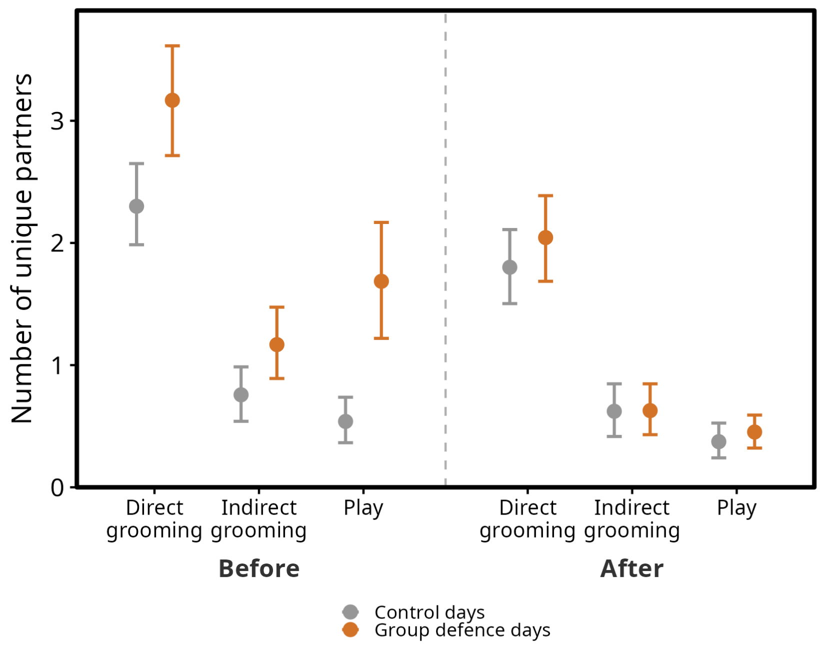

Figure S8: Number of unique affiliation partners (all ages, matched by focal group; left to right: direct grooming, indirect grooming [belonging to the same polyadic grooming cluster without direct contact], play) before and after times of group defence. Points show the mean number of unique partners per observation across group defence days and 1000 resamples of an equal number of matched control days. Error bars are 95% intervals of the estimated mean: for group defence days, non-parametric bootstrap resamples (with replacement) of observations; for control days, across all 1000 iterations.

**Timing of partner access relative to group defence activity (matched by group and focal individual):**

**Independent individuals:**

Table S19: Estimated number of partners by affiliation modality for independent partners.

| Behaviour | Period | Days of group defence | Control days | Mean difference [95% quantiles] | p |
|---|---|---|---|---|---|
| Direct affiliation | Before | 2.78 | 1.92 | 0.86 [0.57, 1.12] | <0.001 |
| Direct affiliation | After | 1.64 | 1.47 | 0.17 [-0.09, 0.42] | 0.218 |
| Direct grooming | Before | 2.43 | 1.84 | 0.59 [0.31, 0.86] | <0.001 |
| Direct grooming | After | 1.55 | 1.41 | 0.13 [-0.11, 0.38] | 0.308 |
| Play | Before | 0.56 | 0.13 | 0.43 [0.34, 0.50] | <0.001 |
| Play | After | 0.09 | 0.08 | 0.01 [-0.04, 0.06] | 0.756 |
| Indirect grooming | Before | 0.91 | 0.59 | 0.33 [0.13, 0.50] | <0.001 |
| Indirect grooming | After | 0.49 | 0.47 | 0.02 [-0.15, 0.18] | 0.788 |

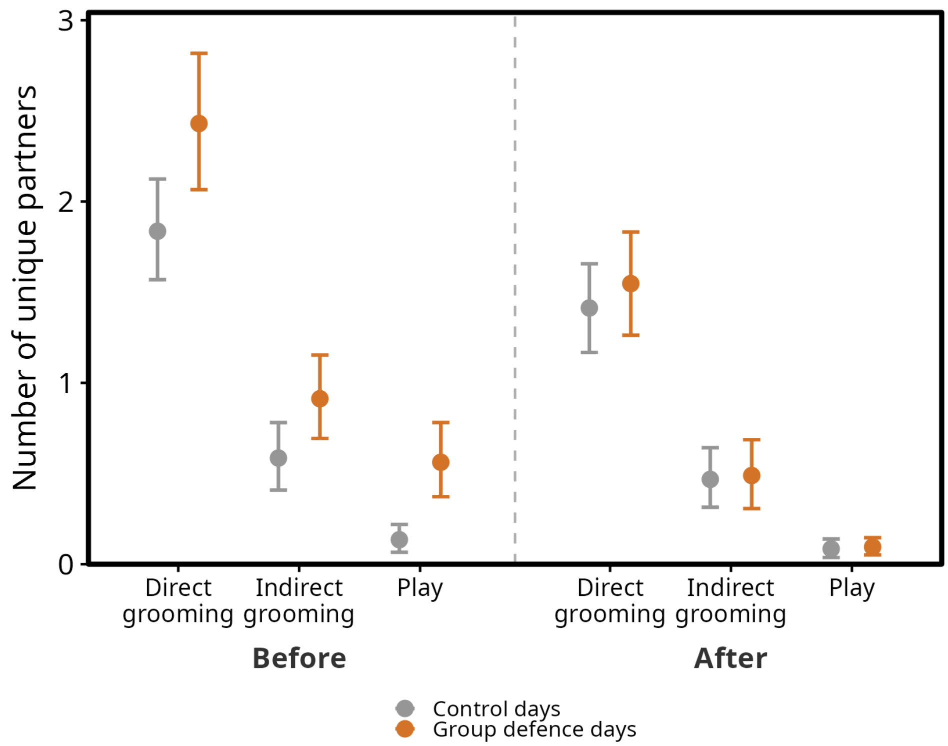

Figure S9: Number of unique affiliation partners (independent-aged, matched by focal group and individual; left to right: direct grooming, indirect grooming [belonging to the same polyadic grooming cluster without direct contact], play) before and after times of group defence. Points show the mean number of unique partners per observation across group defence days and 1000 resamples of an equal number of matched control days. Error bars are 95% intervals of the estimated mean: for group defence days, non-parametric bootstrap resamples (with replacement) of observations; for control days, across all 1000 iterations.

**All ages:**

Table S20: Estimated number of partners by affiliation modality for all partners.

| Behaviour | Period | Days of group defence | Control days | Mean difference [95% quantiles] | p |
|---|---|---|---|---|---|
| Direct affiliation | Before | 4.5 | 2.89 | 1.61 [1.23, 1.96] | <0.001 |
| Direct affiliation | After | 2.37 | 2.19 | 0.18 [-0.17, 0.52] | 0.314 |
| Direct grooming | Before | 3.17 | 2.46 | 0.70 [0.36, 1.01] | <0.001 |
| Direct grooming | After | 2.04 | 1.89 | 0.16 [-0.13, 0.46] | 0.32 |
| Play | Before | 1.69 | 0.59 | 1.09 [0.88, 1.28] | <0.001 |
| Play | After | 0.45 | 0.41 | 0.04 [-0.12, 0.18] | 0.63 |
| Indirect grooming | Before | 1.17 | 0.84 | 0.33 [0.09, 0.55] | 0.01 |
| Indirect grooming | After | 0.63 | 0.68 | -0.05 [-0.29, 0.15] | 0.648 |

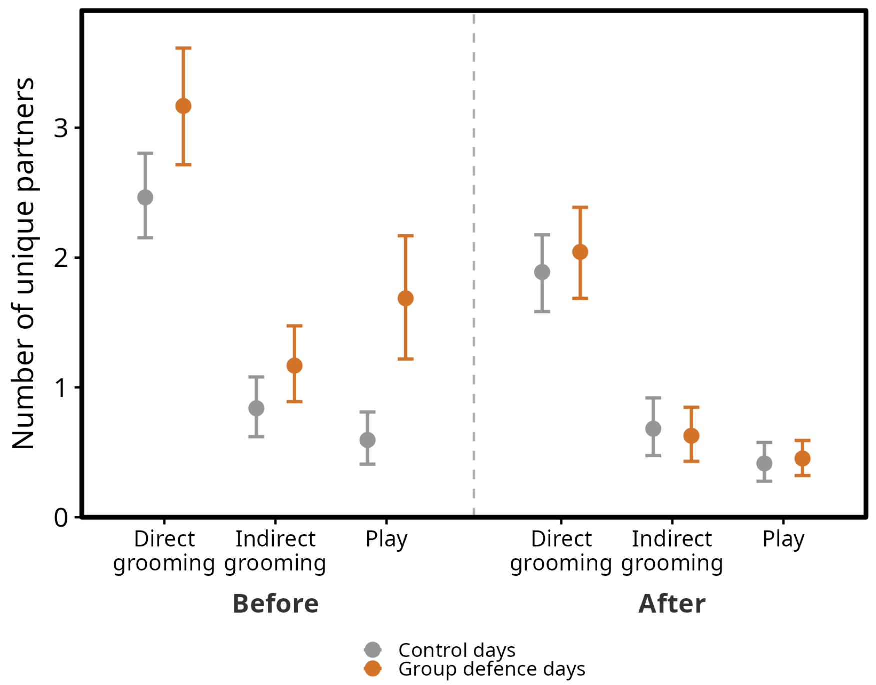

Figure S10: Number of unique affiliation partners (independent-aged, matched by focal group and individual; left to right: direct grooming, indirect grooming [belonging to the same polyadic grooming cluster without direct contact], play) before and after times of group defence. Points show the mean number of unique partners per observation across group defence days and 1000 resamples of an equal number of matched control days. Error bars are 95% intervals of the estimated mean: for group defence days, non-parametric bootstrap resamples (with replacement) of observations; for control days, across all 1000 iterations.
